# WNT signaling coordinately controls mouse limb bud outgrowth and establishment of the digit-interdigit pattern

**DOI:** 10.1101/2024.12.25.629665

**Authors:** Jonas Malkmus, Angela Morabito, Lucille Lopez-Delisle, Laura Avino Esteban, Alexandre Mayran, Aimee Zuniga, James Sharpe, Rolf Zeller, Rushikesh Sheth

## Abstract

Self-organization, such as the emergence of a pattern from a homogenous state, is a fascinating property of biological systems. Early limb bud outgrowth and patterning in mice are controlled by a robust and self-regulatory signaling system, and initiation of the periodic digit-interdigit pattern appears under control of a self-regulatory Turing system. Previous studies established the requirement of WNT and BMP signaling for both early limb bud and digit-interdigit morphogenesis, but the molecular changes underlying the transition from early limb bud signaling to the digit-interdigit patterning system remained unknown. Here, we use small molecule inhibitors to rapidly but transiently block WNT signaling to identify the early transcriptional targets that are altered during disruption and recovery of limb bud and digit development. Together, this study highlights the overarching role of WNT signaling in controlling early limb bud outgrowth and patterning, and establishment of the periodic digit-interdigit pattern. Finally, the transient WNT signaling disruption approach reveals the plasticity and robustness of these self-organizing limb bud and digit patterning systems.

**Summary Statement:** It was unknown how the signaling system controlling early limb bud development transitions to the periodic digit-interdigit patterning system. We show that WNT signaling controls this transition as an integral part of both signalling systems.

## Introduction

The generation of temporally and spatially organized patterns in an embryonic field is regulated by the balance of progenitor cell proliferation and specification. The developing limb bud is a paradigm to study the interplay between growth, patterning, and tissue morphogenesis. Briefly, WNT, BMP, and FGF signaling are first required to establish the dorsal-ventral (DV) limb bud axis and apical ectodermal ridge (AER) during limb bud formation (Geetha-Loganathan et al., 2008; Mariani et al., 2008; Soshnikova et al., 2003; Zeller et al., 2009). In turn, AER-FGF signaling is essential to promote proximal-distal (PD) outgrowth, while the posterior Sonic Hedgehog (SHH) signaling center is required for anterior-posterior (AP) limb bud axis patterning (Zuniga and Zeller, 2020). Concurrent transcriptional activation of the BMP antagonist *Gremlin1* (*Grem1*) by BMP and SHH signaling results in establishment of the robust and self-regulatory SHH/GREM1/AER-FGFs signaling system (Bénazet et al., 2009; Zuniga et al., 1999). This signaling system promotes survival and expansion of limb bud mesenchymal progenitors (LMPs) and the LMP populations that will give rise to the autopod territory and digits (Markman et al., 2023; Michos et al., 2004; Palacio et al., 2024). Around embryonic day E11.25, a gap appears in the distal crescent of the *Sox9* expression domain in mouse forelimb buds, which is the earliest molecular sign of breaking the AP symmetry of the *Sox9* domain (Fig. 1A). This marks the onset of establishing the periodic digit-interdigit pattern as the *Sox9*-expressing cells will give rise to digits while cells that cease *Sox9* expression will give rise to interdigital or soft tissue (Fig. 1A, Akiyama et al., 2005; Bénazet et al., 2012; Raspopovic et al., 2014). Subsequently (∼E11.75), organizing centers termed Phalange Forming Regions (PFRs) are established at the distal tip of the metacarpal primordia, which control the distal growth and extension of digits and phalange formation (Fig. 1A, Fig. S1A,B, Grall et al., 2024; Huang and Mackem, 2022; Montero et al., 2008; Suzuki et al., 2008). While the molecular interactions and gene networks controlling these major morphogenetic processes are well studied, the molecular interactions controlling the transition from the early limb buds signaling system to the digit-interdigit patterning system are unknown.

**Figure 1.**
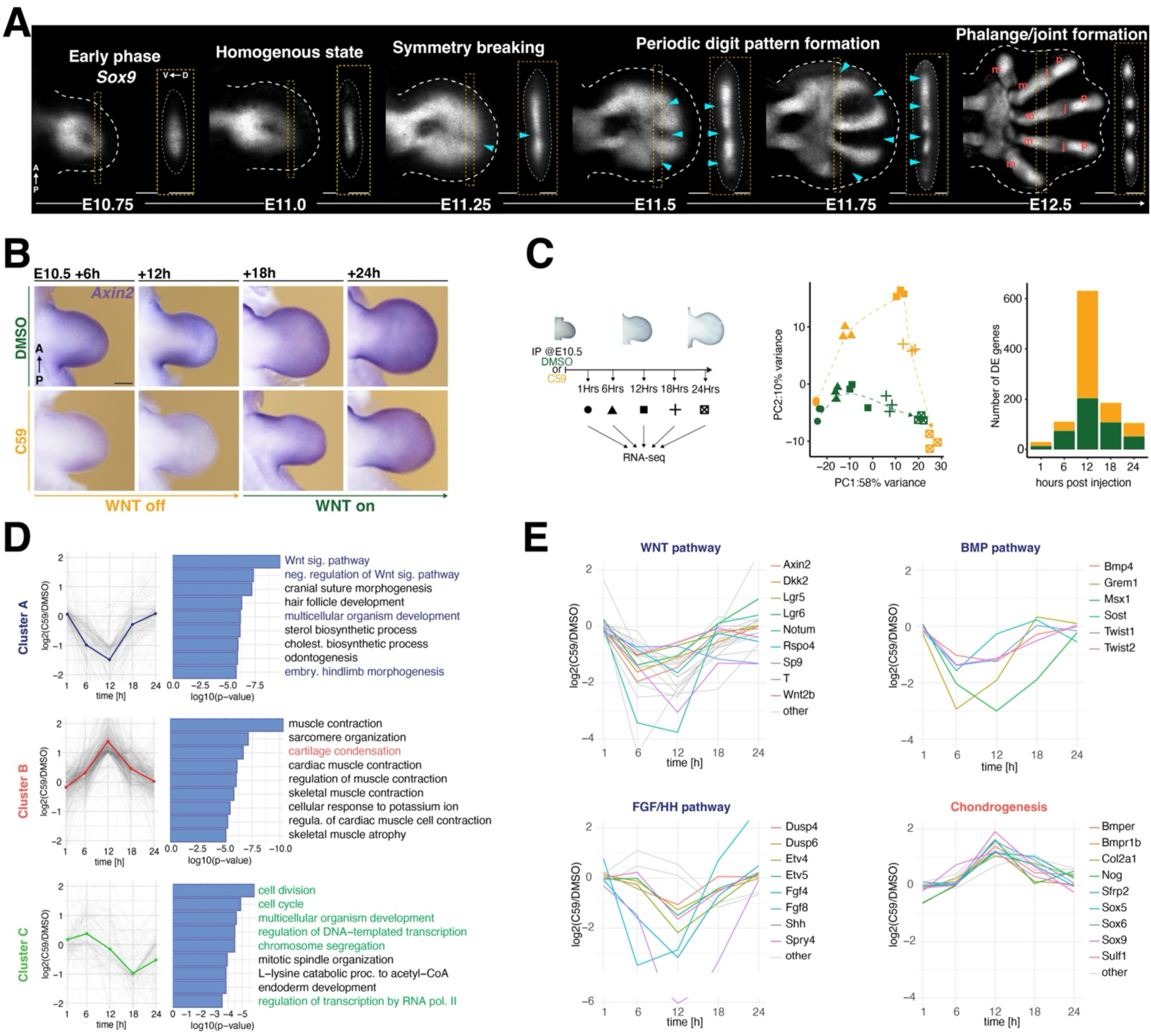
C59-mediated inhibition of WNT signaling leads to transient deregulation of the limb transcriptional programs. **(A)** Whole mount RNA-FISH in wild-type forelimb buds between E10.75 and E12.0 shows the progression of the dynamic spatial distribution of *Sox9* expression from early limb bud outgrowth (E10.75) to late stage digit and phalange development. Notable are the formation of the first interdigit at symmetry breaking (E11.25 and the progressive establishment of periodic digit/metacarpal interdigit pattern. At E12.5 digit 4-2 are composed of three distinct skeletal elements: metacarpals (m), joints (j) and phalanges (p). Interdigits are indicated by cyan arrowheads. A: anterior, P: posterior, D: dorsal, V: ventral. Scalebar: 200µm **(B)** *Axin2* expression detected by WISH in forelimb bud at 6, 12, 18 and 24 hours after injection of either DMSO (control) or Wnt-C59 at ∼E10.5 (n=3 independent biological replicates per stage). Scalebar 250µm. **(C)** At specific timepoints after DMSO (control) or C59 injection into pregnant females (∼E10.5), forelimb buds from embryos were processed for total RNA-sequencing (scheme on the left, n=3 independent biological replicates). PCA analysis shows the differences between the individual timepoints and DMSO (control) and C59-treated samples (middle panel). The bar plots on the right represent numbers of DEGs at each timepoint. Green: control, orange: Wnt-C59 treated samples. **(D)** DEGs were split into three clusters using their profile of fold-changes over time. For each DEG cluster, GO analysis was performed to identify the top 10 enriched biological processes. GO processes most relevant for this study are highlighted by in colour. **(E)** Manually selected DEGS associated with the WNT, BMP and FGF/HH pathway in cluster A (highlighted in blue in panel D) are shown as line plots. Line plots are also shown for DEGs in cluster B that are linked chondrogenic processes (highlighted in red in panel D).

Two morphogen-based hypotheses, positional information and self-organizing reaction-diffusion, have been proposed (Green and Sharpe, 2015; Kondo and Miura, 2010): the positional information hypothesis provides a framework to study specification of AP positional identities in the limb bud mesenchyme by a morphogen gradient, which ultimately endows digits with distinct AP identities (Wolpert, 1969). SHH was identified as the proposed morphogen (Harfe et al., 2004; Riddle et al., 1993), but recent genetic and molecular analysis shows that rather than acting long-range, SHH establishes posterior identities already during onset of limb bud development (Zhu et al., 2008; Zhu et al., 2022). However, the positional information model and the early specification of AP positional identities cannot explain the spacing of the periodic digit-interdigit pattern (Sheth et al., 2012). Based on data-based simulation it has been proposed that digit-interdigit patterning is controlled by self-organizing reaction-diffusion (RD) Turing system (Newman and Frisch, 1979; Raspopovic et al., 2014; Sheth et al., 2012). These simulation led to the proposal that the periodic pattern emerges as a consequence of local interactions among reacting and diffusing proteins, in *senso stricto* not depending on positional information. From Turing patterning systems other than limb buds it is known that interactions of at least two morphogens with different diffusion rates disrupt the homogenous state and produce spatially opposing patterns, which upon reaching steady state manifesting themselves by repeating spots or stripes (Gierer and Meinhardt, 1972; Kondo and Miura, 2010; Turing, 1952). Several studies provide insight into the molecular interactions that are in support of self-organizing Turing mechanisms acting during distal limb bud patterning. For example, molecular analysis support the involvement of Turing-type systems in (a) the initial periodic AP patterning of metacarpals (Raspopovic et al., 2014; Sheth et al., 2012), (b) subsequent reiterative joint formation during PD outgrowth of digit primordia (Grall et al., 2024), and (c) emergence of the fingerprint patterns (Glover et al., 2023).

During limb development, WNT/β-Catenin signaling plays an essential role from initiation of limb bud outgrowth to subsequent establishment of the periodic digit-interdigit pattern. Previous studies have shown that in early limb buds, ectodermal WNT and AER-FGFs maintain the underlying LMPs in an undifferentiated proliferative state and suppress chondrogenesis by restricting *Sox9* expression and BMP activity to the core mesenchyme (Berge et al., 2008; Reinhardt et al., 2019). With respect to digit-interdigit patterning, a Turing model derived from mathematical simulations of experimental data incorporating WNT and BMP signaling and the transcription factor SOX9 predicts that interdigit BMP signaling promotes *Sox9* expression in digit primordia, while SOX9 restricts *Bmp2* expression to the interdigit (Raspopovic et al., 2014). In contrast, WNT signaling and SOX9 are proposed to be mutually antagonistic such that WNT inhibits *Sox9* expression in the interdigit, while SOX9 represses *Wnt* from digit territories. This BMP-SOX9-WNT (BSW) Turing network provides useful predictions for testing the molecular interactions that control the periodic digit-interdigit pattern.

Here, we show that conditional inactivation of *β-Catenin* in mouse limb buds corroborates the requirement of canonical WNT signaling in establishment of the digit-interdigit pattern. However, to overcome the slow kinetics of genetic inactivation and the general disruptive nature of genetic *β-Catenin* inactivation, we resorted to using a small molecule WNT signaling inhibitor. In particular, intraperitoneal (IP) injection of Wnt-C59 results in rapid but transient disruption of WNT signaling, which allowed identification of immediate transcriptional targets of WNT/β-Catenin in early mouse forelimb buds (∼E10.5). This reveals the positive impact of WNT signaling on multiple transcriptional target genes that function as part of the robust SHH/GREM1/AER-FGF signaling system. In addition, this analysis provides insight into the progression from this early limb bud signaling system to the digit-interdigit patterning system. WNT inhibition transiently disrupts establishment of the periodic digit-interdigit pattern, while the expression of genes functioning in chondrogenesis and digit patterning such as *Sox9*, *Sfrp2* and *Sulf1* are upregulated and expanded. During progressive restoration of WNT signaling the periodic digit-interdigit pattern recovers. Gene expression profiling identifies *Wnt2*/*Wnt2a* as ligands expressed in by interdigit mesenchyme and *Sfrp2* and *Sulf1* as two candidate WNT singling modulators that could function in restriction of WNT/β-Catenin signaling to the interdigit mesenchyme. This analysis provides evidence that *Sfrp2* and *Sulf1* could function as part of the Turing type digit-interdigit patterning system during early autopod development.

## Results

### Conditional inactivation of *β-Catenin* reveals the genetic requirement of canonical WNT signaling for digit patterning

The requirement of canonical WNT/β-Catenin signaling for limb bud development was studied using two different genetic approaches. Initially, a tamoxifen-inducible pan-active CRE-transgene (Hayashi and McMahon, 2002) was used for conditional inactivation of *β-Catenin* (Huelsken et al., 2001) during in early limb buds (∼E10.0, *Ctnnb1*^τιc/τιc^, Fig. S2). The expression of *Axin2*, a transcriptional sensor of canonical WNT signaling, is reduced but not completely lost after 48hrs (Fig. S2A) together with genes functioning in the self-regulatory limb bud signaling system that controls limb bud outgrowth and patterning (Fig. S2B,C, Hill et al., 2006). In addition, *Sox9* expression expands into the sub-ectodermal mesenchyme and the presumptive digit territory in *Ctnnb1*^τιc/τιc^ forelimb buds (Fig. S2D), which interferes with establishment of digit-interdigit pattern. Furthermore, *Hoxa13*-CRE-mediated conditional *β-Catenin* inactivation results in downregulation of *Axin2* and expansion of *Sox9* expression in the presumptive digit forming area (Fig. S2E, F, Uenal, 2015), which causes metacarpal fusions and loss of phalanges (Fig. S2G). While this genetic analysis shows that canonical WNT/β-Catenin signaling is required for both the initial digit-interdigit and subsequent phalange pattern; the slow but permanent inactivation do not allow identification of the likely distinct roles WNT signaling during early limb bud outgrowth and digit-interdigit patterning.

### Transcriptional alterations in response to disrupting WNT signaling in mouse limb buds

To circumvent the limitations of conditional genetic inactivation, the efficacy of three different small molecule inhibitors (Chen et al., 2009; Madan et al., 2016; Proffitt et al., 2013) in abrogating WNT signal transduction was assessed by IP injection into pregnant mice at gestational day ∼E10.5 (Fig. S3A, B). This identified Wnt-C59 as the best suited small molecule, which inhibits porcupine activity and WNT ligand secretion (Proffitt et al., 2013) that has been successfully used to inhibit WNT signaling in mouse and Axolotl (Lovely et al., 2022; Szenker-Ravi et al., 2018). A single IP injection of C59 (10μg/gr body weight) at gestational day ∼E10.5 (Fig. S3A, B) causes rapid transcriptional down-regulation of the WNT signaling sensor *Axin2* (left panels, Fig. 1B). Within 6hrs, *Axin2* expression is reduced to low levels but recovers within ∼18 to 24hrs revealing the rapid but transient nature of WNT signal disruption in mouse embryos (right panels, Fig. 1B). To study the impact on gene expression, forelimb buds of DMSO-injected control and C59-treated mouse embryos were collected for RNA-sequencing (RNA-seq) shortly after injection at ∼E10.5 (at 1hr), WNT signal disruption (at 6, 12hrs) and recovery (at 18, 24hrs left panel, Fig. 1C, Table S1). As revealed by PCA analysis, the transcriptional dynamics following inhibition of WNT signaling display distinct temporal trajectories in control and C59-treated forelimb buds (middle panel, Fig. 1C). In both cases, the gene expression profiles separate along the PC1 axis according to developmental time. In particular, the C59-treated forelimb bud samples diverge from their control groups along the PC2 axis between 6-18hrs (middle panel, Fig. 1C). This points to transient transcriptional differences in the gene expression profiles of C59-treated limb buds, which is corroborated by an increase in differentially expressed genes (DEGs) that peaks 12hrs post-C59 injection (right panel, Fig. 1C). Subsequently, the transcriptional differences are reduced and restored close to control levels in C59-treated forelimb buds at 24hrs (middle and right panel, Fig. 1C). Clustering of differentially expressed genes using the profile of the fold-changes over time, reveals three distinct gene clusters with distinct temporal kinetics of transcriptional alterations (Fig. 1D, Fig. S4). In cluster A, DEGs are rapidly downregulated reaching maximum reductions between 6-12hrs, followed by restoration close to control levels at 24 hours. Many DEGs in cluster A function as part of the major signaling pathways required for mouse limb bud outgrowth and patterning (Fig. 1E, Table S2), which in addition to WNT includes the BMP, FGF and SHH pathways (panels WNT, BMP and FGF/SHH, Fig. 1E, Table S3). Conversely, cluster B is enriched in genes that function in chondrogenesis and myogenesis (middle panel, Fig. 1D, Table S1). In particular, DEGs that function in chondrogenesis and/or digit-interdigit patterning such as *Sox9*, *Bmpr1b*, *Sfrp2* and *Sulf1* (Figs. 5, 6 below) are precociously upregulated after Wnt-C59 injection, which reaches its maximum at 12hrs and are reduced again close to control levels by 24hrs (panel Chondrogenesis, Fig. 1E, Fig. S4B, Table S2). Cluster C is enriched in DEGs that function in transcriptional and cell cycle regulation and cell division (bottom panel, Fig. 1D, Table S1). The delayed transcriptional downregulation of DEGs in cluster C peaks at 18hrs and only partially recovers by 24hrs, which indicates that these DEGs are likely altered downstream of disrupting WNT signaling (compare cluster A to cluster C, Fig 1D). The observed transcriptional dichotomy indicates that WNT signaling regulates the balance of LMP proliferation and patterning while inhibiting precocious chondrogenic fates and upregulation of genes function in digit patterning in early limb buds (∼E10.5-11.0).

### Identification of the early transcriptional targets of WNT/β-Catenin signaling in mouse limb buds

C59-mediated WNT inhibition shows that the BMP, FGF and SHH signaling pathways are downregulated with similar rapid kinetics (≤6hrs, Fig. 1E) indicating that these pathways are early targets of WNT signal transduction. Therefore, network analysis using these DEGs was performed to identify the gene regulatory networks whose expression is positively regulated by WNT signaling (Fig. 2A, Table S4). This network analysis pinpoints the primary targets in the WNT, BMP and FGF signaling pathways. To determine the extent to which these alterations may depend on direct regulation by β-Catenin-mediated WNT signaling, specific antibodies for β-Catenin and H3K27ac that marks active enhancer and promoter regions (Andrey et al., 2017), were used for chromatin immunoprecipitation sequencing (ChIP-seq). For both β-Catenin and the histone H3K27ac mark, ChIP-seq of control and C59-treated limb buds was done at E10.5+6hrs (∼E10.75, Fig. 2B). The distribution of β-Catenin close to transcriptional start sites (TSS) and distal region in control limb buds shows that β-Catenin primarily interacts with distal *cis*-regulatory modules (CRMs; control in left panels, Fig. 2B). Motif analysis identifies a TCF/LEF binding sequence as the top enriched motif (Fig. 2C). This is expected as β-Catenin often forms complexes with TCF/LEF1 transcriptional regulators, which mediate the response to canonical WNT signal transduction (Cadigan and Waterman, 2012; Söderholm and Cantù, 2021). Within 6 hours following C59-treatment, β-Catenin binding is depleted both at TSS and distal regions in comparison to control limb buds (C59 in left panels, Fig. 2B). In contrast, the histone H3K27ac profile is not significantly changed (right panels, Fig. 2B). Furthermore, 141 of 197 (72%) of validated VISTA enhancers active in limb buds overlap regions enriched in β-Catenin chromatin complexes (Fig. 2D) (Visel et al., 2007). In addition, the β-Catenin interactions with *cis*-regulatory regions in the genomic landscapes of relevant DEGs were analyzed in control and C59-treated forelimb buds at E10.5+6hrs (Fig. 2E-H and Fig S5). This identifies several β-Catenin binding regions (ChIP-seq peaks) in the *Axin2* genomic landscape that could mediate the rapid transcriptional feedback between *Axin2* expression and WNT/β-Catenin signal transduction in untreated limb buds (profile in black, Fig. 2E). In agreement with the rapid loss of *Axin2* expression following C59 treatment (Fig. 1B), the interaction of β-Catenin complexes with CRMs is depleted (profile in orange at E10.5+6hrs, Fig. 2E). Key transcriptional targets in the BMP pathway that are positively regulated by WNT signaling include the BMP antagonist *Gremlin1* (*Grem1*) and the BMP ligand *Bmp4* (Fig. 2A). Indeed, the limb-specific enhancers in the *Grem1* genomic landscape (Malkmus et al., 2021) and an enhancer regulating *Bmp4* expression in mouse limb buds (Jumlongras et al., 2012) are all enriched in β-Catenin complexes (black profile), while these enrichments are lost within 6hrs after C59 treatment (orange profiles, Fig. 2F,G). This is also the case for three enhancers that regulate *Fgf8* expression in the AER of mouse limb buds (Fig. 2H, Marinić et al., 2013). This analysis shows that these genes are positively and directly regulated by WNT signaling by interaction of β-Catenin complexes with CRM enhancers that regulate their expression in mouse limb buds.

**Figure 2.**
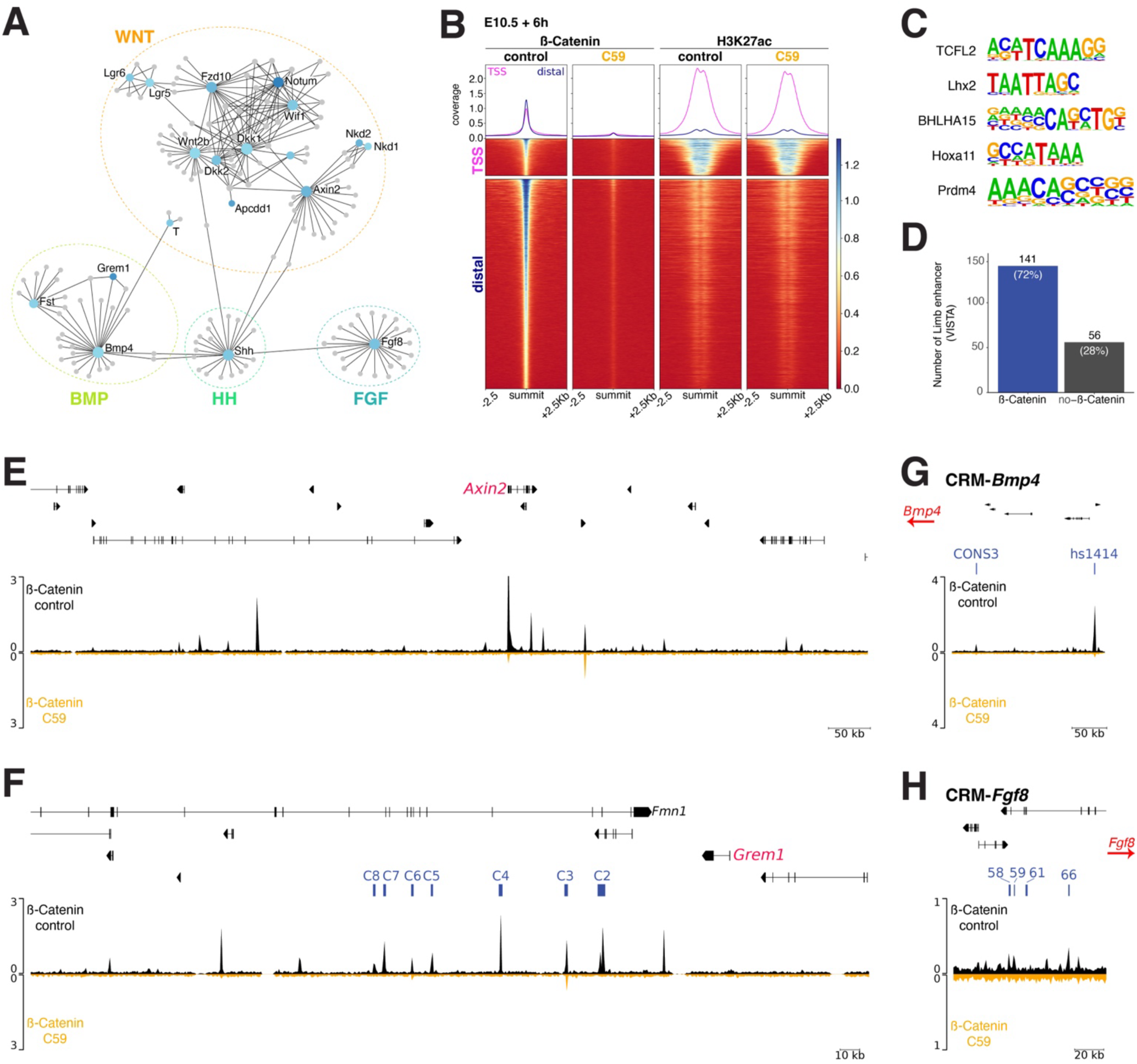
Application of the small molecule inhibitor C59 disrupts WNT signaling. **(A)** Genes downregulated as early as 6hrs post C59 IP injection were used to generate curated gene interaction networks. The subnetwork 1 with 24 seeds (genes) is shown after removal of the *Igf1, Bdnf, Ngf, Tac1* and *Ccl28* seeds. **(B)** ChIP-seq analysis of the β-Catenin interactions with chromatin and the H3K27Ac histone modification profiles in forelimb buds assessed 6hrs after IP injection of DMSO (control) and C59 into pregnant females (∼E10.5, n=2 independent biological replicates). Top: line plots on top show the β-Catenin distribution and H3K27ac profiles at transcriptional start sites (TSS) and distal regions. Heatmaps show the presence or absence of β-Catenin interactions and overlapping H3K27ac modifications in control and C59-treated limb buds. **(C)** The top five *de novo* consensus motifs enriched by β-Catenin ChIP-seq analysis. **(D)** Validated enhancers with reporter activity in limb buds were intersected with β-Catenin ChIP-seq peaks. 141 of 197 VISTA limb bud enhancers encode regions enriched in β-Catenin chromatin complexes. **(E)** For the *Axin2* locus (exons shown as black bars on top), the β-Catenin ChIP-seq profile (E10.5+6hrs) is plotted for control (black profile) and C59-treated limb buds (orange profile). Inhibition of WNT signaling results in loss of most β-Catenin ChIP-seq peaks. **(F)** The *Grem1 cis*-regulatory landscape encodes seven CRMs (C2-C8, indicated in blue) that are required for the dynamic regulation of *Grem1* expression in mouse limb buds (exons indicated on top). All seven CRMs encode regions enriched in β-Catenin chromatin complexes (black profile). These ChIP-seq peaks are reduced or lost upon inhibition of WNT signaling (orange profile). **(G)** CRM-*Bmp4*: CONS3 is a validated enhancer regulating *Bmp4* expression and VISTA hs1414 is an enhancer with AER-specific activity. Both CRMs (blue) encode regions enriched in β-Catenin ChIP-seq peaks (black profile) that are lost when WNT signaling is inhibited (orange profile). **(H)** CRM-*Fgf8*: Four CRMs (blue) in the *Fgf8* genomic landscape are active in the AER. Three of them are marked by β-Catenin ChIP-seq peaks (black profile) that are lost upon inhibiting WNT signaling (orange profile).

### WNT signaling is required for the self-regulatory signaling system that controls limb bud outgrowth and patterning

The analysis thus far establishes that WNT signaling positively regulates key genes in the self-regulatory limb bud signaling system that interlinks mesenchymal SHH (Fig. S5E) with BMP4/GREM1 and AER-FGF signaling (Fig. 2A, 2F-H) to coordinately control AP axis pattering and PD progression of limb bud outgrowth. To assess the impact of transient WNT signal disruption on the spatial dynamics of gene expression, mouse *Shh*, *Grem1* and *Fgf8* HCR probes were multiplexed for fluorescent whole mount RNA *in situ* hybridization (RNA-FISH, Fig. 3A). Analyzing their expression in the same forelimb buds reveals the rapid reduction of *Grem1* transcripts and *Fgf8* expression in the AER (6-12hrs), which is followed by progressive recovery after 18-24hrs (∼E11.0-E11.5, Fig. 3A). This downregulation is paralleled by almost complete loss of AER-FGF signal transduction as reflected by reduction of the expression of the FGF sensor *Dusp6* from the distal limb bud mesenchyme (∼E11.0-E11.5, Fig. 3B) (Kawakami et al., 2003). The transient disruption not only lowers *Grem1* transcripts and protein (Fig. 3A, Fig. S6A) but also mesenchymal *Bmp4* expression (Fig. 3C). The reduction and recovery of BMP activity is apparent from the spatio-temporal kinetics of pSMAD1,5,9 activities (pSMAD, Fig. 3D) and dynamic expression of BMP target genes in C59-treated limb buds (*Msx2*, *Id1*, Fig. S6B,C). Posterior *Shh* expression is reduced, but recovers to a lesser extent (Fig. 3A), which is reflected by the progressively reduced spatial expression of its transcriptional target *Ptch1* (Fig. S6D). Following transient inhibition, the limb bud signaling system scales down by 6hrs, but largely recovers by 24hrs (Fig. 3). In all cases, the developing autopods of C59-treated embryos are smaller (Fig. 3, Fig. S6), which substantiates the WNT requirement for LMP cell proliferation that was uncovered by the initial DEG analysis (cluster C, Fig. 1D). This analysis establishes the WNT signaling pathway as an integral part of the self-regulatory signaling system that controls early limb bud outgrowth and expansion of the LMP populations that will give rise to the autopod primordia (Desanlis et al., 2020; Markman et al., 2023; Palacio et al., 2024; Sheth et al., 2013).

**Figure 3.**
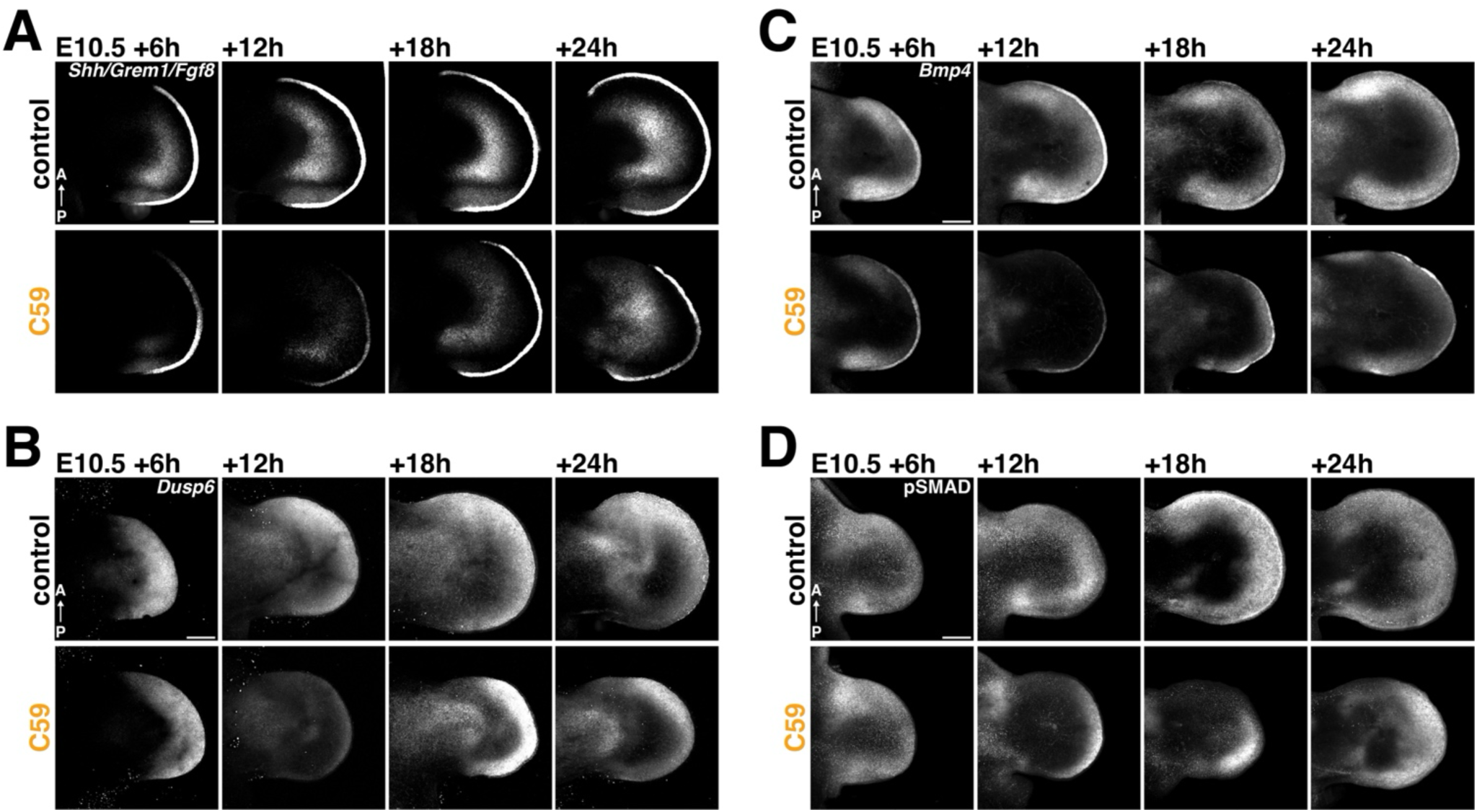
Spatio-temporal response of the self-regulatory SHH/GREM1/AER-FGF signaling system to inhibition of WNT signaling. **(A-C)** RNA-FISH analysis at four timepoints (E10.5+6, +12, +24hrs) following DMSO (control) or C59-injection into pregnant females reveals the spatio-temporal dynamics of gene expression during transient disruption and recovery of WNT signaling. **(A)** *Shh* (posterior mesenchyme), *Grem1* (posterior-distal) and AER-*Fgf8* expression assessed in the same forelimb buds using the same hairpin amplifier for all three probes. **(B)** Expression of the FGF-transcriptional target *Dusp6* in the same forelimb buds as shown in panel A. **(C)** Spatial dynamics of *Bmp4* expression in control and C59-treated forelimb buds. For all limb buds in panels A-C the entire Z-stack range is shown as maximum intensity projection. **(D)** Whole mount immunostaining of the pSMAD1,5,9 (pSMAD) distribution in limb buds reveals active BMP signal transduction. A selected Z-stack range (midsection) is shown as maximum intensity projections. n=3 independent biological replicates were analysed for all time points shown in all panels A-D. A=anterior, P=posterior. Scalebars: 200µm.

### WNT inhibition during early limb bud outgrowth also disrupts subsequent digit-interdigit patterning

Genetic analysis underscores the essential role of β-Catenin-mediated WNT signal transduction in the digit-interdigit patterning system (Fig. S3). C59-mediated transient disruption of WNT signaling at E10.5 also allows to assess the effects on digit ray patterning, as the progressive recovery of WNT signaling (Fig. 1B-E) overlaps the time window of symmetry breaking (∼E11.25, Fig. 1A) and establishment of the periodic digit-interdigit pattern (≥E11.5, Fig. 1A). To gain insight into the effects on pattern formation, *Sox9* expression was analyzed in control and C59-treated limb buds between 12-36hrs after injection (∼E10.5, Fig. 4A). In contrast to appearance of first digit-interdigit separation in control forelimb buds (indicated by red and blue arrowheads, upper panels Fig. 4A), the *Sox9* expression domain is distally expanded and there is no evidence for the onset of digit-interdigit patterning at the equivalent stage in C59-treated forelimb buds (E10.5+12hrs, lower left panel Fig. 4A). This is consistent with negative regulation of *Sox9* expression by WNT signaling (Fig. S2)(Hill et al., 2005). During recovery of WNT signaling (Fig 1B, C), *Sox9* expression is restricted again from the distal-most mesenchyme and a periodic digit-interdigit pattern is established by E10.5+36hrs (∼E11.5-12.0, lower middle and right panels Fig. 4A). The *BMP receptor1b* (*Bmpr1b*) is expressed with temporal dynamics similar to *Sox*9 during distal progression of limb bud development (upper panels Fig. S7A, see also Fig. 1E). Following C59-treatment, *Bmpr1b* expression is expanded into the distal-most mesenchyme at E10.5+12hrs (∼E11.0, lower left panel Fig. S7A). During recovery from WNT inhibition, *Bmpr1b* expression is progressively re-restricted to the forming digit primordia by 24-36hrs (∼E11.5-E12.0, lower middle and right panels Fig. S7A). Conversely, expression of the *Bmp2* is restricted to the interdigit mesenchyme during digit-interdigit patterning (upper panels Fig. 4B) and its expression in the distal mesenchyme is downregulated when WNT signaling is disrupted (lower panels Fig. 4B). During recovery of WNT signaling, *Bmp2* expression is progressively reestablished (at +24 hrs) and restricted to interdigit at 36hrs (∼E11.5-12.0, lower right panel Fig. 4B). In addition to *Bmp2*, the crescent-shaped *Grem1* expression domain in the dorsal and ventral mesenchyme (upper left panel Fig. 4C) gets restricted to the forming interdigit mesenchyme during digit-interdigit patterning (upper middle and right panels Fig. 4C). Following C59 treatment, *Grem1* expression is lost (lower left panel Fig. 4C), but as expression recovers it is restricted to the interdigit mesenchyme (lower middle and right panels Fig. 4C). These results show that WNT signaling positively regulates the expression of the BMP ligand *Bmp2* and the BMP antagonist *Grem1* in the distal and interdigit mesenchyme. Thirty-six hours after C59-treatment, the autopod primordia remains smaller (lower right panels, Fig 4A-C) and only to 3 or 4 digits (86%) form as a consequence of the spatially constraint digit territory (Fig 4D). This reduction in digit numbers agrees with previous studies that linked autopodial progenitor cell proliferation to the number of digits formed (Alberch and Gale, 1983; Lopez-Rios et al., 2012).

**Figure 4.**
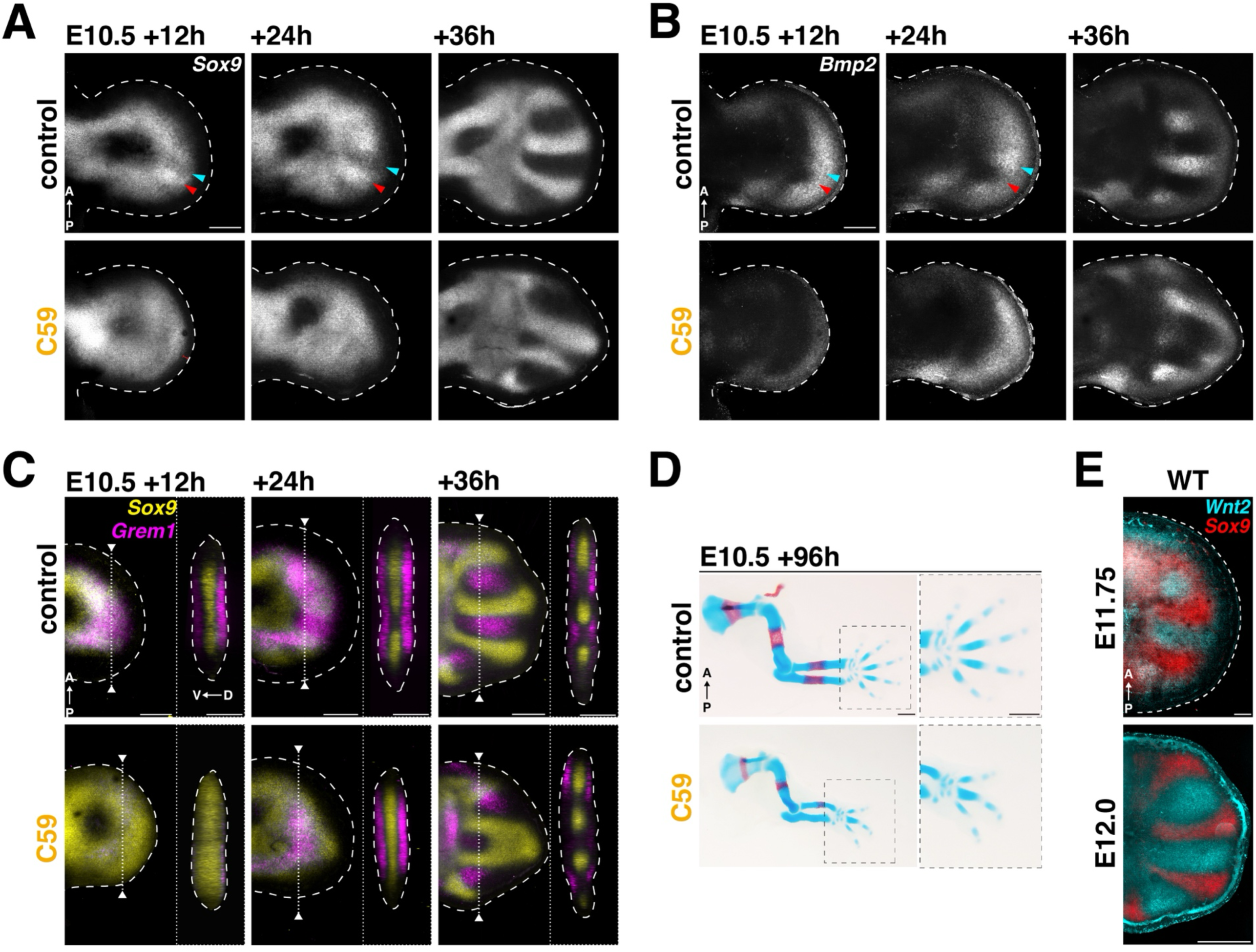
Destabilization and reestablishment of the digit pattern following transient disruption of WNT signaling. RNA-FISH analysis of the expression dynamics following DMSO (control) or C59-injection into pregnant females (∼E10.5). The 3 timepoints illustrate the disruption (+12hrs) and recovery phase (+24, +36hrs). **(A)** Spatio-temporal kinetics of *Sox9* expression. **(B)** Spatio-temporal kinetics of *Bmp2* expression. **(C)** *Sox9* (yellow) and *Grem1* (magenta) expression dynamics. Right panels show virtual cross-sections at the level indicated by dotted lines in the left panels. **(D)** Alizarin red (bone)/Alcian blue (cartilage) staining of E14.5 limb skeletons of embryos treated with DMSO (control) or C59 at E10.5. Disrupting WNT signaling leads to oligodactyly in all cases, in most cases (52%) 3 digits remain. Others display four (39%), two (6%) or one (3%) digit(s). n= 31 biological replicates. A=anterior, P=posterior, D=dorsal, V=ventral. Scalebar: 500µm. **(E)** Colocalizations of *Sox9* (digit primordia, red) with *Wnt2* (interdigit mesenchyme cyan, RNA-FISH) at the indicated stages. For the panels in A-C and D a selected z-stack range is shown as maximum intensity projection. Scalebars for A-C, E: 200µm.

To date the source and identity of the WNT ligand(s) functioning in the periodic digit-interdigit patterning system has remained elusive. However, C59-mediated inhibition of WNT signaling identifies the *Wnt2b* ligand among the DEGs downregulated in the WNT pathway (Fig 1E). This is likely relevant to establishment of the periodic digit pattern as *Wnt2b* and its paralogue *Wnt2* are expressed by the interdigit mesenchyme in wildtype forelimb buds (Fig. S7B)(Witte et al., 2009), which indicates that the interdigit is a source for WNT ligand(s) functioning in digit-interdigit patterning (Fig. 4D). This is consistent with the WNT signaling activity being restricted to the interdigit mesenchyme during digit patterning either by (a) SOX9 suppressing *Wnt* ligand expression and/or (b) inhibition of WNT activity in the *Sox9*-expressing digit territory by negative modulation of signal transduction and/or WNT antagonists (Raspopovic et al., 2014). Therefore, *Sox9-*positive cells are a likely source of negative regulators acting in phase with *Sox9* upon WNT signaling disruption. Indeed, gene expression profiling identifies *Sulf1*, an extracellular modulator of WNT (and BMP) signaling and the extracellular WNT antagonist *Sfrp2* that transiently upregulated together with *Sox9* following C59-mediated inhibition of WNT signaling (panel Chondrogenesis Fig. 1E, Fellgett et al., 2015; Kawano and Kypta, 2003; Otsuki et al., 2010; Sahota and Dhoot, 2009; Satoh et al., 2006). RNA-FISH shows that *Sox9*, *Sulf1* (Fig. 5A) and *Sfrp2* (Fig. 5C) are expressed in distinct spatial domains during wildtype autopod development (control, Fig. 5A, C). During digit-interdigit patterning, *Sulf1* expression is restricted to the emerging *Sox9-*positive digit primordia (middle panel Fig. 5A) and its expression extends distally during digit ray elongation (right panel Fig. 5A). Conversely, *Sfrp2* expression is restricted to the interdigit (middle panel Fig. 5C) with a distinct proximal expression boundary at the presumptive wrist territory (right panel Fig. 5C). Inhibition of WNT signaling causes distal-anterior and dorso-ventral expansion of *Sox9* and *Sulf1* expression at E10.5+12hrs (∼E11.0, left panel Fig. 5B). Concurrently, *Sfrp2* expression expands precociously anterior within the distal limb bud mesenchyme (left panel Fig. 5D, compare to middle panel, Fig. 5C). During WNT recovery, the digit-interdigit periodicity of the *Sox9*, *Sulf1* and *Sfrp2* expression domains is re-established (right panels, Fig. 5B, D). These results show that inhibiting WNT signaling disrupts the patterning process as an expansion of *Sox9* expression at the expense of interdigit gene expression is observed. However, this is reversed and digit-interdigit patterning resumes as WNT signaling recovers, which points plasticity and/or robustness of digit/interdigit patterning even if the digit territory and number of digits formed are reduced following the transient inhibition of WNT signaling.

**Figure 5.**
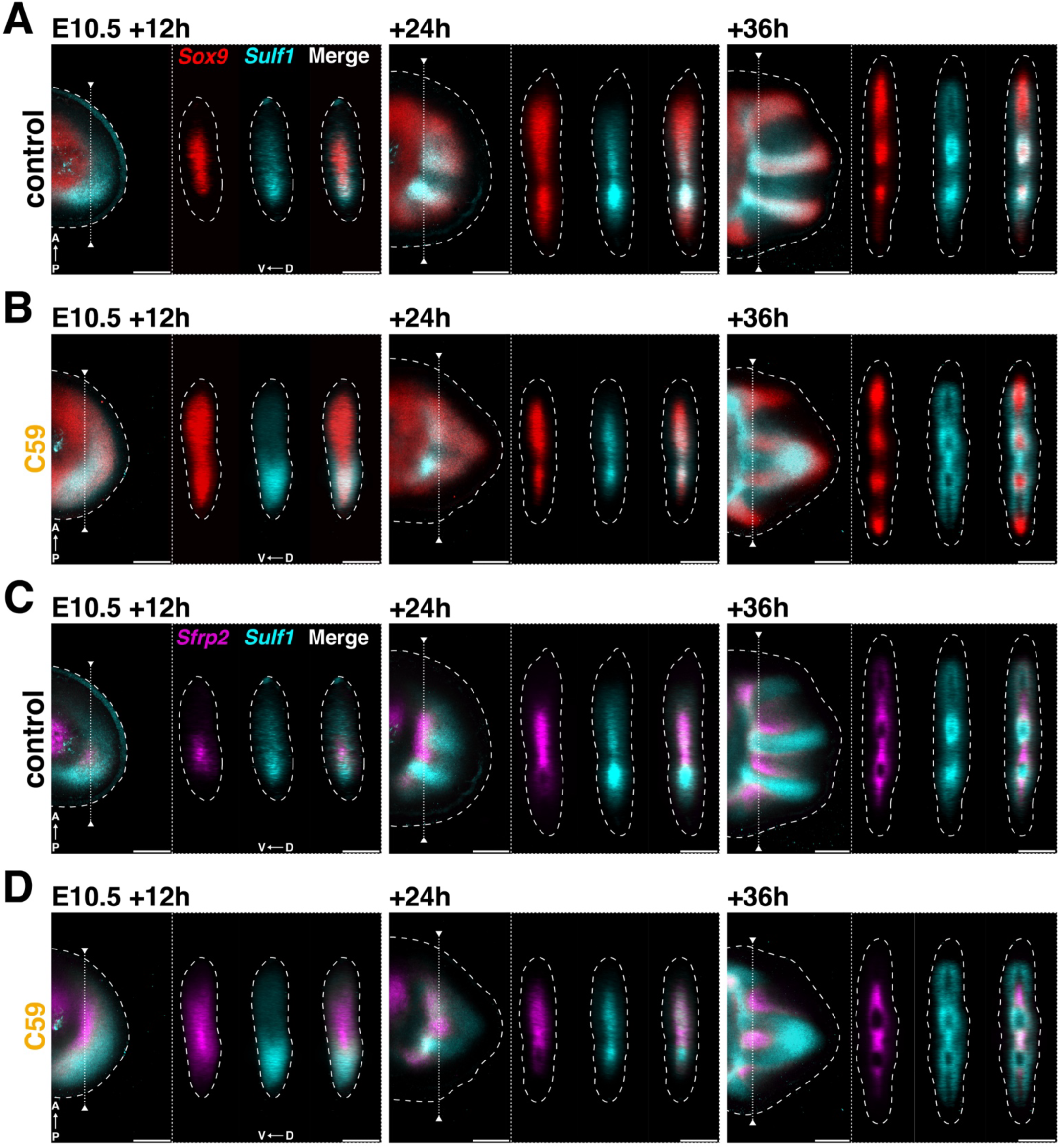
The periodic expression of the signaling modulator *Sulf1* and the antagonist *Sfrp2* requires WNT signaling. **(A-B)** Colocalization of *Sox9* (red) with *Sulf1* (cyan) shows the spatio-temporal dynamics in control (panels in A) and C59 treated (panels in B) forelimb buds between 12-36hrs after injection (∼E10.5). Left panels: representative virtual mid-sections of the 3D images are shown. Right panels: show virtual cross-sections of the dorso-ventral axis at the level indicated by a line in the left panels. **(C, D)** Colocalization of *Sfrp2* (red) with *Sulf1* (cyan) in the limb buds shown in panels A, B. A=anterior, P=posterior, D=dorsal, V=ventral. n=3 independent biological replicates for all timepoints. Scalebars: 200µm.

### Transient inhibition of WNT signal during the establishment of the periodic digit-interdigit pattern

To gain insight into the spatiotemporal dynamics of digit-interdigit patterning during the progression of digit rays (metacarpals), WNT signaling was disrupted at ∼E11.5. The observed downregulation and recovery of *Axin2* expression (Fig. S8A) establishes that the kinetics of transiently disrupting WNT signaling at this later stage is comparable to C59-treatment at early forelimb bud stages (∼E10.5, Fig. 1B). C59 was injected at ∼E11.5, as inhibiting WNT signaling at this stage overlaps the period of AP digit patterning (Fig 1A, 6A), but precedes the onset digit phalange patterning in mouse forelimb buds that is controlled by the PFR signaling center (≥E12, Fig. 1A, Fig. S1B, Huang and Mackem, 2022). C59 IP injection at ∼E11.5 causes expansion and upregulation of *Sox9*, *Bmpr1b* and *Sulf1* expression into the interdigit territories and distal-most mesenchyme compared to control forelimb buds at E11.5+12hrs (∼E12.0, left panels Fig. 6A-C). As WNT signaling is restored by E11.5+ 24hrs (Fig. S8A), the spatial expression of *Sox9*, *Bmpr1b* and *Sulf1* is again restricted to the digit ray primordia (∼E12.5, right panels Fig. 6A-C). However, aberrant digit curvatures (assessed by *Sox9* and *Bmpr1*b expression, Fig. 6A-D) and proximal bifurcations or broadening of the metacarpal base of the anterior digit 2 are also observed (asterisks Fig. 6, n=8/9). In addition *Sox9* is also expressed as thin stripes in the interdigit territory (arrowhead Figure 6A, n=3/6). Effects on interdigit gene expression were assessed by analyzing the spatial distribution of *Sfrp*2 and *Bmp2* in control and C59-treated forelimb buds (Fig. 6D, Figure S8B). In C59-treated forelimb buds, *Sfrp2* expression expands precociously distal within the interdigit, but is excluded from the distally expanded *Sox9* domain at E11.5+12hrs (∼E12.0, left panels Fig. 6D). In comparison to controls, *Bmp2* expression is lost from the distal mesenchyme but maintained in the proximal interdigit territory (∼E12.0, left panels Fig. S8B). Both *Bmp2* and *Sfrp2* expression recover as WNT signaling is restored at E11.5+24hrs (∼E12.5, right panels Fig. 6D, Fig. S6B). Interestingly, the altered interdigit gene expression patterns are complementary to the metacarpal alterations at this stage (Fig. 6D). However, all these alterations seem transient as the final digit/metacarpal morphology and pentadactyl state are preserved (n=18/19, lower panel, Fig. 6E).

**Figure 6.**
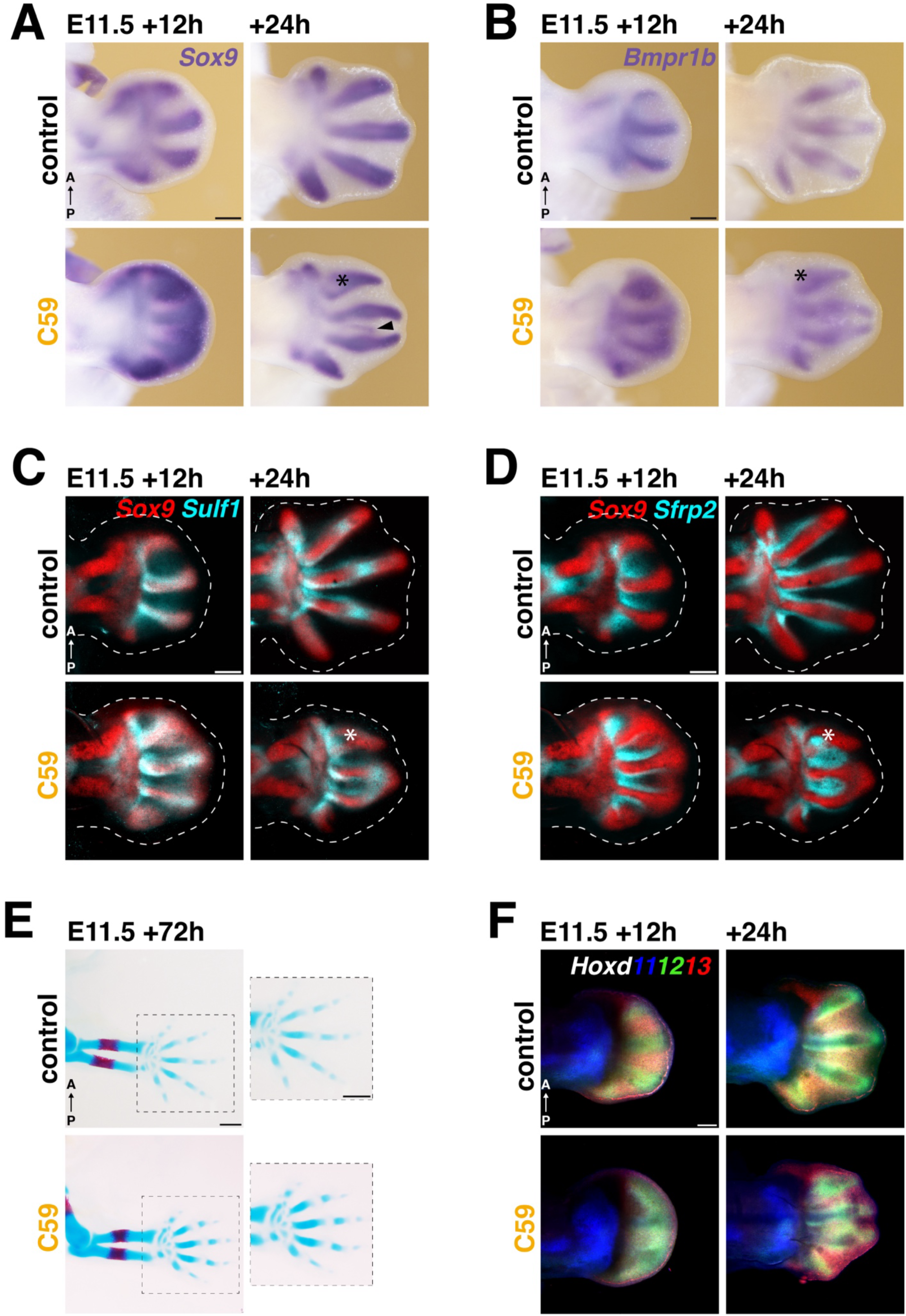
Alteration of the digit and interdigit gene expression patterns following WNT inhibition during early digit ray development. Forelimb buds were analysed 12 and 24hrs after DMSO (control) and C59 IP injection at ∼E11.5. **(A, B)** The *Sox9* and *Bmpr1b* expression dynamics in control and C59-treated forelimb buds. **(C)** Colocalization of *Sox9* (red) with *Sulf1* (cyan) in control and C59-treated limb buds. **(D)** Colocalization of *Sox9* (red) with *Sfrp2* (cyan) in the same limb buds shown in panel C. For the limb buds shown in panels C and D selected z-stack ranges are shown.. Scalebars in panels A, B: 250μm; C,D: 200μm. **(E)** Limb skeletons of embryos exposed to DMSO (control) or C59 at ∼E11.5. Bone is visualized by Alizarin red and cartilage by Alcian blue. Transient disruption of WNT signaling does not alter pentadactyly (n=18/19). Scalebar: 500μm. All limb buds in panels A-F are oriented with anterior (A) to the top and posterior (P) to the bottom. **(F)** Colocalization of *Hoxd11* (blue), *Hoxd12* (green) and *Hoxd13* expression in control and C59-treated forelimb buds. For limb buds in panel E, the entire Z-stack ranges are shown as maximum projections. Scalebars: 200μm. n=3 independent biological replicates were analysed for all genes and timepoints shown in panels A-D,F

Distal Hox genes have been proposed to modulate the digit wavelength as part of the BSW Turing digit-interdigit patterning system (Raspopovic et al., 2014; Sheth et al., 2012). RNA-FISH analysis of *Hoxd11, Hoxd12* and *Hoxd13* in control forelimb buds reveals that *Hoxd12* expression is highest in digit primordia, while *Hoxd13* expression is highest in interdigits at E12.0-E12.5 (top panels, Fig. 6F, Fig. S8C: individual patterns). Following C59-treatment at ∼E11.5, *Hoxd12* expression is expanded into the interdigit mesenchyme, while *Hoxd13* expression is reduced (E11.5+12hrs, bottom left panel, Fig. 6F, left panels Fig. S8D: individual patterns). After restoration of WNT signaling, the distinct digit-interdigit expression domains of *Hoxd12* and *Hoxd13* recover to similar levels as control limb buds (E11.5+12hrs, right panels Fig. 6F, Fig. S8C compare to Fig. S8D). Taken together, this analysis shows that C59-mediated transient inhibition of WNT signaling during early limb bud outgrowth and later during ongoing digit-interdigit patterning significantly impacts the expression of key WNT targets genes as part of the early limb bud signaling system (Bénazet et al., 2009) and periodic digit-interdigit patterning (Raspopovic et al., 2014; Sheth et al., 2012). As WNT signaling recovers, the disrupted gene expression patterns and digit development recover largely, which points to significant robustness and/or plasticity of the limb bud and digit-interdigit patterning systems.

## Discussion

In this study, we use IP injection of the small molecule inhibitor C59 into pregnant females to rapidly and transiently disrupt WNT signaling during mouse limb bud outgrowth and digit patterning. This analysis establishes that (1) WNT/β-Catenin signaling is required for maintenance and propagation of the SHH/GREM1/AER-FGF self-regulatory signaling system; (2) the coordination of growth, restriction of *Sox9*-expressing progenitors to the chondrogenic core mesenchyme and developing digit rays; and (3) initial establishment of the periodic digit-interdigit pattern during early autopod development. In particular, inhibition of WNT signaling during forelimb bud outgrowth and patterning (∼E10.5) results in rapid downregulation of target genes in the SHH, BMP and FGF signaling pathways, which establishes the WNT pathway is an integral part of the self-regulatory limb bud signaling system that controls limb bud outgrowth and patterning (Bénazet et al., 2009; Zuniga and Zeller, 2020). Previously validated CRMs in the genomic landscapes of *Shh* (Lettice et al., 2003), *Bmp4* and *Grem1* (Jumlongras et al., 2012; Malkmus et al., 2021), and AER-*Fgf8* (Marinić et al., 2013) are enriched in β-Catenin chromatin complexes, pointing to regulation by WNT/β-catenin signal transduction. This is corroborated by β-Catenin depletion from CRMs following inhibition, which occurs with kinetics similar to downregulation of target gene expression. Canonical WNT target gene activation depends on β-Catenin binding to cofactors, such as TCF/LEF and the assembly of a large multiprotein complexes at *cis*-regulatory elements (Cadigan and Waterman, 2012; Söderholm and Cantù, 2021). Interaction of β-Catenin with these multiprotein complexes at WNT-responsive elements promotes transcription (Söderholm and Cantù, 2021). It is likely that the C59-mediated WNT signaling disruption and decline of nuclear complexes affects the enhancer-promoter interactions while multiprotein complexes including TCF/LEF and other cofactors remain bound to *cis*-regulatory elements. In such a poised state, restoration of WNT signaling would allow rapid re-establishment of enhancer-promoter interactions and target gene transcription. This is corroborated by the kinetics of transcriptional changes observed by gene expression profiling and spatial-temporal gene expression analysis during WNT-signaling disruption and recovery (this study). For example, the rapid loss and recovery of *Grem1* transcripts and proteins following C59-treatment underscores the sensitivity of the dynamic spatio-temporal transcriptional regulation by WNT/β-catenin signaling. The recovery from early C59-mediated inhibition of WNT signaling also covers the period during which the self-regulatory limb bud signaling system transitions to periodic digit-interdigit patterning system. This transition is paralleled by genes required for autopod development becoming independent of SHH (Panman et al., 2006), and the progressive shut-down of the SHH/GREM1/AER-FGF signaling system (Farin et al., 2013; Verheyden and Sun, 2008). During this transition, interactions between the WNT, BMP signaling and the *Sox9* transcription factor have been proposed to initiate the periodic digit-interdigit pattern system with characteristics of a self-organizing Turing mechanism (Raspopovic et al., 2014). Both conditional genetic inactivation of β-Catenin during autopod development and C59-mediated WNT inhibition at ∼E10.5 and ∼E11.5 causes expansion of the *Sox9* expression domain and prevents or perturbs the digit-interdigit pattern (this study). Although C59-mediated inhibition of WNT signaling blocks both canonical and non-canonical WNT signaling (Proffitt et al., 2013), the genetic analysis reveals the requirement of β-Catenin, i.e. canonical WNT signaling for limb bud and digit development (this study). This is corroborated by a study that showed that the initial digit-interdigit pattern is established correctly when the non-canonical *Wnt5a* ligand is inactivated (Yamaguchi et al., 1999). These and other studies indicate that canonical WNT/β-Catenin signal transduction is required for establishment of the initial digit-interdigit pattern, while non-canonical WNT signaling functions in regulating growth and elongation of digit primordia as part of a convergent-extension mechanism (Parada et al., 2022).

It has been shown that a fundamental property of Turing systems is their ability to self-reorganize after disruption, often resulting in curved stripes and loops. This is associated with the intrinsic self-organizing behavior of Turing systems, whereby the positions and spacing of the periodic pattern arises spontaneously and is maintained over space. Experimental perturbation in different developmental systems have shown how Turing patterns self-reorganize the pigmentation stripes in fish (Kondo and Asai, 1995) and the rodent fur stripe pattern (Johnson et al., 2023), rugae formation during palate patterning (Economou et al., 2012) and avian tracheal cartilage ring patterning (Kingsley et al., 2023). In the present study, transient inhibition of WNT signaling allowed us to evaluate the self-reorganization potential of the periodic digit-interdigit pattern in mouse embryonic limb buds. Perturbation of WNT signaling at E10.5 prevents establishment of initial periodicity of *Sox9* expression; however, upon restoration, a periodic pattern emerges, which resulting in variable curved digit morphologies as observed when disrupting a Turing system (see above). Transient inhibition of WNT signaling at ∼E11.5 results in (a) expansion of *Sox9* expression into the interdigit territory, which might transiently perturb the interdigit pattern and b) together with changes in digit and interdigit gene expression patterns causes transient bifurcations of digit gene expression domains, ectopic *Sox9* stripes and curved morphologies. These alterations of digit-interdigit gene expression and the subsequent recovery of their periodic patterns point to the substantial plasticity of digit-interdigit patterning system by the proposed Turing mechanism (Raspopovic et al., 2014; Sheth et al., 2012). Following establishment of the initial digit-interdigit pattern, cartilage condensation and distal elongation of digit primordia are controlled by *Wnt5* dependent convergent-extension movements that couple PFR formation to digit elongation and phalange specification (Parada et al., 2022). Finally, recent analysis of phalange pattern formation in chicken and mouse limb buds shows that establishment of the periodic joint and phalanx pattern is controlled by a Turing mechanism involving the BMP ligand GDF5 and the BMP antagonist Noggin (Grall et al., 2024).

Several WNT ligands, WNT antagonists, and signaling modulators are expressed by the limb bud mesenchyme and ectoderm during establishment of the digit-interdigit pattern (Geetha-Loganathan et al., 2008; Murphy et al., 2022; Witte et al., 2009). It has been suggested that WNT signaling is required for symmetry breaking (Raspopovic et al., 2014). In agreement, this study shows that the *Wnt2*/*Wnt2b* ligands and the WNT antagonist *Sfrp2* are expressed in the forming interdigit territories, while the signaling modulator *Sulf1* is expressed in the *Sox9* positive mesenchyme (Fig. 7B-E). These expression patterns permitted simulation of a variety of network topologies using the RDNets tool (Marcon et al., 2016) and stability analysis showed that a simple circuit incorporating WNT2, β-Catenin and SFRP2 can function as a classical activator-inhibitor Turing-type circuit (Fig 7B). It is possible that this circuit is initiated during symmetry breaking to function in establishment of the interdigit pattern. Furthermore, negative modulation of WNT signaling by SULF1 downstream of SOX9 would likely contribute to restrict WNT activity to the interdigit territory (Fig. 7C, D). The BSW Turing model predicts that BMP signaling from the interdigit mesenchyme promotes *Sox9* expression in the mesenchyme that will give rise to digit territories. In addition, it has been shown that SULF1 positively modulates BMP signaling during chondrogenesis (Meyers et al., 2013; Ratzka et al., 2008). Therefore, integrating SULF1 into results a Turing circuit with substrate-depletion rather than activator-inhibitor network topology (Fig. 7E). Finally, combining these two additional Turing circuits generates a Turing pattern in which the signal modulator SULF1 and antagonist SFRP2 are expressed out-of-phase in digit and interdigit territories, respectively, to modulate and restrict WNT signaling (Fig. 7E). Previous genetic analysis of *Sfrp1* and *Sfrp2* in mouse embryos revealed functional redundancy, but inactivation of both *Sfrp* genes causes a pleiotropic phenotype that includes stunting and broadening of limb skeletal elements and preaxial polydactyly (Satoh et al., 2006). Similarly, *Sulf1* and *Sulf2* regulate several developmental signaling pathways by cleaving sulfate groups from heparan sulfate proteoglycans (Ai et al., 2003; Fellgett et al., 2015; Otsuki et al., 2010; Sahota and Dhoot, 2009). Genetic loss-of-function analysis reveals their requirement in modulating endochondral bone differentiation (Ratzka et al., 2008), but this not informative with respect to the proposed role *Sulf1* in digit-interdigit pattern. It is known that signaling modulators contribute to the systems’ robustness; and here we evaluated *Sfrp2* and *Sulf1* based on their spatial and temporal expression kinetics during mouse limb bud and digit patterning. In summary, the present study based on transient inhibition of WNT signaling reveals the substantial plasticity of the periodic digit-interdigit patterning system. The present study together with the consecutive Turing mechanism that controls phalange-joint segmentation indicates that spatial modulation of signaling by antagonists contributes both robustness and evolutionary plasticity (Grall et al., 2024).

**Figure 7.**
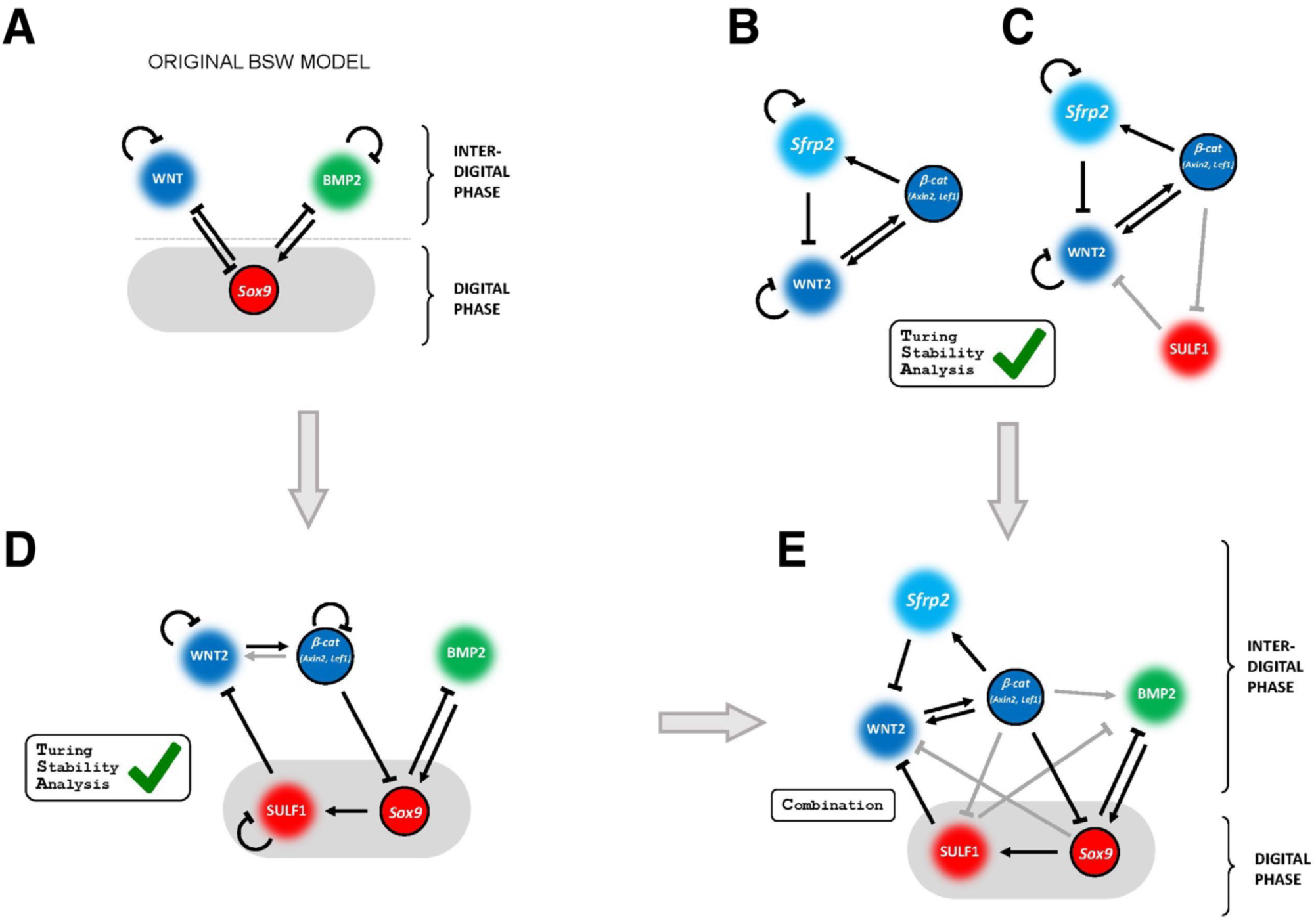
Evaluation of WNT2 and WNT as potential signaling modulators in the three node BSW Turing network. **(A)** scheme of the BSW Turing model proposed by Raspopovic et al (2014). **(B)** Simple model simulations of 3-in-phase node circuit that incorporates WNT2 and SFRP2. The simulations indicate that this circuit can act as a classical activator-inhibitor module. **(C)** Extension of the 3 node network shown in panel B to incorporate the WNT modulator SULF1 expressed in the digit mesenchyme (out-of-phase with SFRP2) as a negative modulator of WNT2. **(D)** Extension of the original BSW model (panel A), to include SULF1 downstream of *Sox9* to repress the diffusible WNT2 ligand the digit mesenchyme (in grey). This model also includes β-Catenin that acts in a cell-autonomous Manner. Mathematical simulations using by RDNets shows that this extended BSW network functions as a substrate-depletion Turing system (rather than activator-inhibitor model). **(E)** Numerical simulations of the 4-node Turing circuit (panel C) in combination with the 5-node circuit (panel D) show that this extended network can function as a Turing circuit.

## Materials and Methods

### Animals

#### Ethics statement and approval of all animal experimentation

All studies involving mice were performed in accordance with national laws and approved by the national and local regulatory authorities as mandated by law in Switzerland. All proposed animal studies were approved by the Regional Commission on Animal Experimentation and the Cantonal Veterinary Office of Basel (national license 1951) in accordance with Swiss laws and the 3R principles.

#### Mouse strains and embryos

In line with the refine and reduce 3R principles, all strains were bred into a Swiss Albino (*Mus musculus*) background as only robust phenotypes manifest in this strain background and the numbers of embryos and litter sizes are large (≥12-15 embryos per pregnant females). Embryos of both sexes at the developmental ages indicated were used for experimental analysis. For conditional *β-Catenin* inactivation, TMCRE*Ctnnb1*^τιc/+^ mice (tamoxifen-inducible CRE) (Hayashi and McMahon, 2002) were crossed to *Ctnnb1*^τιc/τιc^ mice (Catnnb1lox mice – B6.129-Ctnnb1^tm2Kem^/KnwJ, Huelsken et al., 2001). Tamoxifen injections: the tamoxifen stock solution (Sigma-Aldrich T5648) was prepared by dissolving tamoxifen powder in corn oil (Merck C8267) at a concentration of 20mg/ml. Per mouse (35-40gr), one dose of 5mg TM in 250μl corn oil solution was IP injected to induce conditional gene inactivation.

#### Small molecule inhibition of WNT signaling

Wnt-C59 (Tocris, 5148-10mg), a potent porcupine inhibitor that blocks secretion of WNT ligands (Proffitt et al., 2013; Szenker-Ravi et al., 2018), was prepared as a 5mg/ml stock solution in DMSO (Sigma-Aldrich, D8418-250ml). Prior to injection, the C59-stock solution was diluted in vehicle solution (1xPBS, 0.5% methylcellulose (Sigma-Aldrich, M0262-100G), 0.01% Tween-20 (Sigma-Aldrich, 93773-1kg) and 5% DMSO, Szenker-Ravi et al., 2018) to the required working concentration. For DMSO (control) and C59 injections, pregnant Swiss Albino females were weighed in the morning of gestational days 10 or 11 prior to timed IP injection. The C59 (experimental) and DMSO (control) injection volume, respectively was calculated in relation to bodyweight. For controls, 10 % DMSO (in 1xPBS) was IP injected in a volume of 10μl per gram of bodyweight. To inhibit WNT signaling; 10μg/10μl C59 per gram mouse was IP injected at the desired time points (see results).

#### Embryo collection and staging

For whole mount in situ hybridization, RNA-FISH or immunostaining embryos were collected in ice cold PBS and subsequently fixed in 4% PFA overnight. After washing in PBS embryos were dehydrated stepwise into 100% MEOH and stored at least 1 day or longer at −20°C. Wildtype and control injected limb buds were staged by somite counting and limb bud shape. Due to the requirement of WNT signaling for somitogenesis C59-treated embryos (and following conditional β-Catenin inactivation) mutant limb buds were staged (1) according to the time elapsed at the time of isolation since C59 or tamoxifen IP injection; and (2) by comparing limb bud sizes and shapes to wildtype and control limb buds.

#### Limb skeletal preparations

Embryos were collected in the late afternoon of embryonic day 14, washed in 1xPBS and fixed in 95% ethanol at room temperature (RT) overnight. Following fixation, embryos were stained for 24 hours in 0.03% (w/v) Alcian blue (Sigma-Aldrich, A3157) diluted in 80% ethanol and 20% glacial acetic acid (Sigma-Aldrich, 100063). Then were washed for 24hrs in 95% ethanol. Subsequently, embryos were pre-cleared for 30 minutes in 1% KOH (Sigma-Aldrich, 105033) and counterstained with 0.005% (w/v) Alizarin (Sigma-Aldrich, A5533) in 1% (w/v) KOH. Then, embryos were cleared in glycerol/1% KOH by stepwise increases in the glycerol concentration (20%, 40%, 60%, 80% glycerol) and preserved in 80% glycerol in water. Alcian blue detects cartilage, while alizarin red stains ossified bone. Analysis was conducted on n>3 embryos per genotype.

#### Whole mount RNA *in situ* hybridization using BM-Purple staining

A standard protocol for whole-mount RNA in situ hybridization was used (Haramis et al., 1995). Following rehydration, embryos underwent bleaching in 6% hydrogen peroxide that was followed by digestion with 10 μg/ml proteinase K (duration adjusted based on embryonic stage and adapted for genes expressed by the limb bud ectoderm). Following prehybridization at 65°C for at least 3hrs, embryos were incubated overnight at 70°C in hybridization solution containing 0.2-1μg/ml heat-denatured antisense riboprobe to detect the transcripts of interest. The next day, embryos were extensively washed, and non-hybridized riboprobe digested with 20μg/ml RNase at 37°C for 45min. After additional washes and pre-blocking, embryos were incubated overnight at 4°C with anti-digoxigenin antibody (1:5000, Roche 11093274910). Following several washes to remove excess antibodies, RNA-riboprobe hybrids were visualized by incubation in BM Purple staining solution (Roche 11442074001). Development of the BM Purple signal stopped during exponential phase before reaching saturation.

#### Fluorescent whole mount RNA in situ hybridization: HCR-RNA-FISH

The HCR RNA FISH technology (Molecular Instruments) was used in combination with the improved HCR RNA-FISH step-by step protocol that reduces autofluorescence by embryonic tissues and blood vessels (Morabito et al., 2023). Briefly, rehydrated and stage matched limb buds where subjected to photochemical bleaching (2x 30min, at RT) followed by incubation in detergent solution for two hours at 37°C. To detect gene expression, limb buds were hybridized with specific mouse HCR probes (Molecular Instruments for 16hrs (or overnight). In case of low level expression (e.g *Wnt2a*) the HCR probe concentration of was doubled and the incubation time extended to 20-24hrs. Signal amplification was done as described in the step-by step protocol, but for difficult to detect transcripts, the amplifier concentration was doubled and the incubation time increased to 24hrs. During washing in 5xSSCT, nuclei were counterstained with DAPI (1:1000, Sigma-Aldrich, D9542). In order to wash away the high salt concentration, limb buds were washed in 1xPBS for 2min and then immersed in the fructose-glycerol clearing solution. Samples were cleared for minimally 24hrs prior to imaging.

#### Whole mount immunofluorescence

Rehydrated age-matched limb buds were briefly washed in PBS, then permeabilized and unspecific binding blocked in 5xSSCT, 1% Triton and 10x donkey serum for ≥3hrs with gentle shaking at RT. The following primary antibodies were used: goat-anti-GREM1 (1:50, R&D Systems, AF956), rabbit-anti-pSMAD1.5.9 (1:200, Cell Signaling Technology, 13820S). Limb buds were incubated with the respective antibodies diluted in 5xSSCT, 1% Triton and 10x donkey serum for 72hrs at RT with gentle shaking. The limb buds were washed in 5xSSCT, 1% Triton and 10X donkey serum for 4hrs with several changes. Depending on the multiplexed primary antibodies, individual or a combination of the following secondary antibodies were used: donkey anti-rabbit-488 (1:250, Jackson,711-545-152), donkey anti-rabbit-555 (1:250, Invitrogen, A31572), donkey anti-rabbit 647 (1:250, Invitrogen, A31573) and anti-goat 647 (1:250, Invitrogen, A21447). Limb buds were incubated with secondary antibodies diluted in 5xSSCT, 1% Triton and 10x donkey serum containing DAPI (1:1000) at RT with gentle shaking. After washing in 5xSSCT, 1% Triton and 10x donkey serum limb buds were quickly washed in 1xPBS and the refractory index adjusted as described for above for RNA-FISH.

#### Image acquisition and data processing

Images were acquired using a confocal spinning disc microscope with 10x objective (10x/0.45 CFI Plan Apo), a confocal spinning disc scan unit (Yokogawa Spinning Disk CSU-W1-T2) and a Nikon Ti-E, Hamamtsu Flash 4.0 V2 CMOS camera. Images intended for 2D maximum intensity projections were acquired with a 5μm Z-step size, while for 3D reconstruction purposes step size was reduced to 1μm. For 2D maximum projections raw images were processed using FIJI. The entire or a specific stack-range was selected and stacked (“Z Project…”, “start=X stop=Y projection=[Max Intensity]”). As autofluorescence was much reduced by the photochemical bleaching prior to fluorescent detection of gene expression patterns, only brightness and contrast adjustments were made for RNA-FISH analyses. Three-dimensional images constructed from 2D image stacks were generated using the IMARIS software. Therefore, raw files were converted to the .ims format and brightness and contrast values adjusted. Virtual sections were obtained from 3D images using either the “Section” or “Clipping Plane” tools.

#### RNA-sequencing time course

Stage matched limb buds from DMSO or C59-treated embryos were fine-dissected and washed in fresh ice-cold PBS; immediately transferred to RNA-later (Sigma-Aldrich R0901) and stored overnight at −4°C. The next day, samples were transferred to −80°C for long-term storage. RNA isolation was performed using the RNeasy Kit (Qiagen 74104) following manufacture instructions. After performing QC analysis on a Fragment Analyzer (Advanced Analytical Technologies, Inc.), RNA was stored at −80 until shipping to the company Novogene (UK). Standard Illumina library preparations and sequencing was performed by Novogene. After quality control libraries were sequenced as PE150 on a NovaSeq6000.

#### ChIP-seq

Dissected forelimb bud tissues were crosslinked using 1% formaldehyde/1xPBS at RT for 12min and crosslinking was stopped in 125mM glycine solution. For β-Catenin ChIP an initial crosslinking step was performed using DSG (disuccinimidyl glutarate, Sigma-Aldrich # 80424) for 40 min at RT as described (Sheth et al., 2016). After lysis in hypertonic buffer, chromatin fragments were generated by sonication. Immunoprecipitation was performed at 4°C overnight using polyclonal anti-β-Catenin antibodies (Invitrogen 71-2700, 5μg per sample) or anti-histone H3(acetyl K27) antibodies (ChIP Grade Abcam ab4729, 5μg per sample). The chromatin complexes were immunoprecipitated using magnetic beads (Fisher Scientific 11202D). After washing beads in RIPA buffer, DNA was eluted from beads followed by overnight reverse cross-linking. Libraries were generated using the next-generation library preparation kit from Takara Bio (Japan) according to manufacturer instructions and the ChIP-Seq libraries were sequenced using a NextSeq instrument (Illumina).

#### Bioinformatics analysis

The calculations have been performed using the facilities of the Scientific IT and Application Support Center of EPFL.

#### RNA-seq analysis

Low quality bases and Truseq adapters were removed with Cutadapt v1.16 (Martin, 2011) using the following sequences:

-a GATCGGAAGAGCACACGTCTGAACTCCAGTCAC
-A GATCGGAAGAGCGTCGTGTAGGGAAAGAGTGTAGATCTCGGTGGTCGCCGTATCATT -q 30-m 15

Reads were mapped on mouse genome assembly mm10 with STAR (Dobin et al., 2012) version 2.7.0e using ENCODE parameters and custom gtf (https://doi.org/10.5281/zenodo.7510406). Counts and coverage were computed at the same time. The values for <u>F</u>ragments <u>p</u>er <u>k</u>ilobase of transcript per <u>m</u>illion mapped reads (FPKM) were computed using the cufflinks version 2.2.1 program (Trapnell et al., 2010) using the following parameters:

--max-bundle-length 10000000--multi-read-correct --library-type “fr-unstranded” -b
/home/ldelisle/genomes/fasta/mm10.fa --no-effective-length-correction -M MTmouse.gtf -G custom.gtf.

Average coverage between replicates was computed using the bigwigAverage from deepTools version 3.5.5. FPKM values excluding genes on chromosomes X, Y and M were transformed with log2 (1+FPKM). PCA was computed (centered, non-scale) using a selection of 2000 genes with the highest variance. The same matrix was used to compute spearman correlation between samples and clustered with the ward.D2 method. Counts from STAR were restricted for protein coding genes and genes on chromosomes X, Y and M were excluded. Differential analysis was performed with DESeq2 version 1.34.0 (Love et al., 2014) on R 4.1.3 for each experimental timepoint after comparing the 3 control and the 3 C59-treated replicates. DEGs with an adjusted p-value below 0.05 and an absolute log2 fold-change above 1 were selected. All genes significantly altered minimally at one timepoint between control and C59-treated limb buds were considered for clustering. The log2 fold-change values estimated by DESeq2 were centered, scaled (per gene) and used to compute the Pearson’s correlation between genes. Genes were clustered using the ward.D2 method and the tree was cut into 3 clusters. The log2 fold-changes displayed in Figure 2 are DESeq2 log2 fold-change (before scale transformation). GO analysis was performed using GOseq version 1.56.0 (Young et al., 2010) including the genes of each cluster and using all protein coding genes as background.

#### ChIP-seq analysis

Low quality bases and Truseq adapters were removed with Cutadapt v1.4 (Martin, 2011) using the following sequences:

-a GATCGGAAGAGCACACGTCTGAACTCCAGTCAC
-A GATCGGAAGAGCGTCGTGTAGGGAAAGAGTGT -q 30 -m 15)

Filtered reads were mapped with Bowtie2 version 2.4.5 (Langmead and Salzberg, 2012) with default parameters. Alignments with MAPQ below 30 were filtered out with samtools version 1.16.1(Danecek et al., 2021). Peaks were called and coverage was performed with macs2 version 2.2.7.1 (Zhang et al., 2008) using the following parameters:

-g 1870000000 --call-summits --format BAMPE -B, input was not used.

Coverage was normalized by the million fragments after filtering. Average coverage between replicates was computed with bigwigAverage from deepTools version 3.5.5 (Ramírez et al., 2016). Consensus peaks were obtained as follows: 1. peaks were called using all replicates with equal contributions as follows. Duplicates were removed from the BAM files of each replicate with Picard version 2.27.4 (http://broadinstitute.github.io/picard/). Each BAM was subsampled to identify the number of reads of the smallest BAM. Peaks were called using all subsampled BAM with the same criteria as for individual replicates. 2. Only peaks with overlapping summits in both replicates were kept. In particular, individual narrowPeaks were overlapped using BEDTools multiinter version 2.30.0 (Quinlan and Hall, 2010) and only intervals present in both replicates were kept. The narrowPeak file of step 1 was intersected with the output of multiinter and was filtered in order to keep only narrowPeaks with summits overlapping the output of multiinter. Consensus β-Catenin ChIP-seq summits in control conditions were filtered to remove summits overlapping the ENCODE blacklist (Amemiya et al., 2019). Any summit located less than 1kb from a transcriptional start site (TSS) of the custom GTF (see RNA-seq analysis) was considered a TSS. Others were considered as distal peaks. Heatmaps centered on consensus β-Catenin summits at TSS or distal regions were generated using deeptools version 3.5.5 (Ramírez et al., 2016) with a bin size of 10bp at -/+3.5kb. These summits were used to identify de novo transcription factor binding motifs in a 200bp window using Homer version 4.11 (Heinz et al., 2010). VISTA enhancers coordinates were downloaded from https://enhancer.lbl.gov/ and lifted over from mm9 to mm10 and intersected with the consensus β-Catenin summits.

#### Gene Network analysis

A protein-protein interaction network was generated from 73 genes downregulated at 6hrs after C59 IP injection (Dataset S4) using the online tool “NetworkAnalyst” (Zhou et al., 2019). The following settings were used: String database, confidence score 850, no experimental evidence, First order-network. The Subnetwork1 included 24 interconnected seeds of which *Tac1, Ccl28, Igf1, Bdnf* and *Ngf* (not relevant for this study) were removed for representation purposes.

#### Turing network simulations

To explore whether hypothetical regulatory networks were capable of creating Turing patterns we used two approaches. For Figure 7, we used the public online tool RDNets (Marcon et al., 2016), which allows the user to graphically specify network topologies, and performs mathematical analysis to determine if a network is compatible with the Turing conditions. Topologies can be fully specified (both the sign of regulatory links, and whether each node is diffusible or not), or ambiguities can be allowed and the program will explore different options. Due to its complexity, for the full combined network in panel H, we performed numerical simulations in a spatial domain using the modelling platform LimbNET (Matyjaszkiewicz and Sharpe, 2024) instead of an analytical solution. We manually found suitable parameter values that successfully created a Turing pattern.

### Dataset and code availability

The RNA- and ChIP-sequencing datasets generated for this study are available under GEO GSE275784. The published CC dataset was downloaded from GEO GSE84793 (Andrey et al., 2017) and converted from mouse genome assembly mm9 to mm10 using the liftover tool (https://genome-store.ucsc.edu/). Tracks were generated with the pyGenomeTrack tool (Lopez-Delisle et al., 2020) and input commands are deposited on Github including the codes used for NGS sequence analysis: https://github.com/lldelisle/scriptsForMalkmusEtAl2024

## Acknowledgements

The authors would like to thank A. Offinger and her team for outstanding mouse husbandry. We would like to thank P. Lorentz from the DBM Microscopy Core Facility for technical support throughout the project. We are grateful to D. Duboule (EPFL) for providing support and access to the EPFL Scientific IT and Application Support Center for the bioinformatics analysis. We are grateful to Erkan Uenal for performing the *Hoxa13*-Cre conditional *Ctnnb1* inactivation analysis. We would like to thank Luciano Marcon for useful discussions about the Turing circuits and the use of RDNets.

## Author contributions

R.S. conceived and supervised the study in discussion and with input from A.Z. and R.Z, who acquired the necessary funding. R.S. J.M. and A.Mo. performed all experimental analysis. L.L.D. and A.Ma. performed all bioinformatics analysis for the RNA- and ChIP-sequencing datasets. L.A.E. and J.S. performed all mathematical calculations and model simulations for the different possible Turing network topologies. Figures were prepared and the manuscript written by R.S., J.M., R.Z. and J.S. (Turing model simulations) with input from all authors into the manuscript draft.

## Supplementary information

contains eight figures and four tables.

