## Supplementary Figures and Legends for "WNT signaling coordinately controls mouse limb bud outgrowth and establishment of the digit-interdigit pattern"

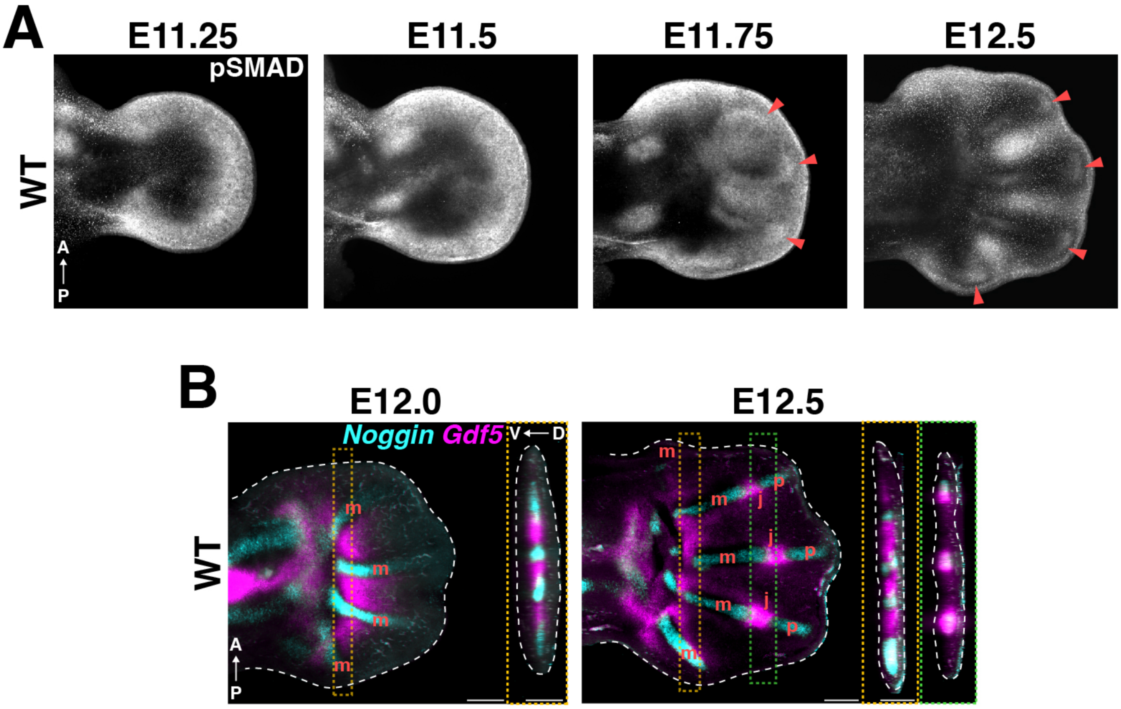


**Fig. S1. The Phalange forming region become molecularly apparent ~E11.75 7and the first joint and phalange have formed by E12.5 in forelimbs.** (A) Developmental series of the pSMAD1,5,9 distribution by wholemount immunofluorescence in wildtype forelimb buds between E11.25 and E12.5. Red arrow heads point to the appearance of the PFR pattern. By E11.75, crescents of high pSMAD expression in the distal tip of forming digit rays reveal the presence of PFRs that become well separated at E12.5. (B) Whole mount RNA-Fish of *Noggin* (cyan) and *Gdf5* (magenta) in wildtype limb buds show that up to E12.0 metacarpals (m) are forming but the future joint marker *Gdf5* is expressed in the proximal part of the interdigit domains. By E12.5, *Gdf5* is also expressed in developing joints (j), which divide the *Noggin*$^{+}$ digit rays into metacarpals (m) and the first phalanges (p). RNA-FISH and immunofluorescence images are shown as maximum intensity projection of a selected Z-stack range. n=3 independent developmental timeseries were analyzed for panels A and B. A=anterior, P=posterior, D=dorsal, V=ventral.

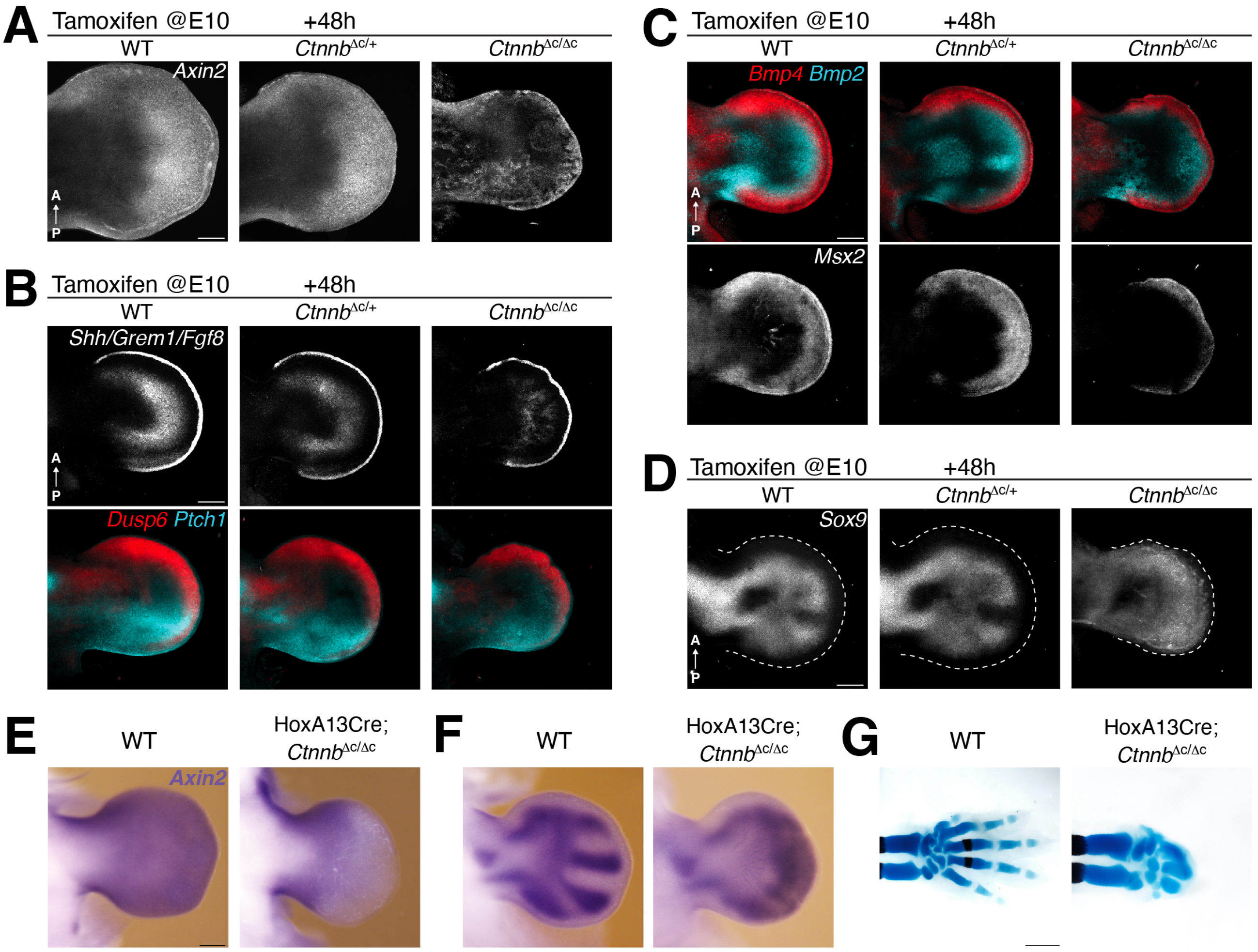


**Fig. S2. Temporal kinetics of conditional genetic β-Catenin gene inactivation in mouse forelimb buds.** (A-D) The expression of key genes functioning in growth, patterning and chondrogenesis were assessed by RNA-FISH in wildtype (Wt), heterozygous (*Ctnnb*$^{\Delta c/+}$) and homozygous (*Ctnnb*$^{\Delta c/\Delta c}$) forelimb buds at 48hrs after tamoxifen injection at ~E10. (A) Expression of the WNT transcriptional sensor *Axin2* is significantly reduced but not lost at E10+48hrs. (B) Upper panels: *Shh*, *Grem1* and AER-*Fgf8* expression were detected in the same channel. While *Shh* expression is significantly reduced/lost, *Grem1* expression is patchy and AER-*Fgf8* expression is somewhat reduced. Lower panels: in the same forelimb buds, the transcriptional sensors *Ptch1* (cyan, for SHH) and *Dusp6* (red, for AER-FGF signaling) were assessed. (C) Upper panels: *Bmp2* (cyan)and *Bmp4* (red) expression in forelimb buds. Both *Bmp2* and *Bmp4* expression are reduced but not lost in *Ctnnb*$^{\Delta c/\Delta c}$ forelimb buds. Lower panels: in the same forelimb buds, *Msx2* expression was assessed. *Msx2* expression is reduced and distalized in *Ctnnb*$^{\Delta c/\Delta c}$ forelimb buds. (D) The *Sox9*

expression domain is significantly expanded in *Ctnnb*$^{\Delta c/\Delta c}$ forelimb buds. n=3 independent biological replicates were analysed for all genotypes and stages. All images are shown as maximum intensity projection of the entire Z-stack range. Forelimb buds are oriented with anterior to the top and posterior to the bottom. Scale bars: 200µm. (E) *HoxA13*Cre-mediated *β-Catenin* inactivation in the distal limb bud mesenchyme. In comparison to wildtype controls, *Axin2* is downregulated in the developing handplate (reminiscent of the *HoxA13* expression domain. (F) *Sox9* is expressed by the distinct digit ray progenitors in wildtype forelimb buds (left panel). In the *βCatenin* deficient hand plates, the characteristic *Sox9* expression in the periodic digit domains is absent as *Sox9* expression is expanded in the distal mesenchyme (right panels). (G) Comparison of the developing wildtype(left ) and *HoxA13*Cre *β-Catenin* deficient autopod skeleton (right). In mutant forelimbs only 2-3 malformed and distally fused digits have formed at E14.5. n≥3 independent biological replicates were analysed for all probes and developmental stages. Limb buds are oriented with anterior to the top and posterior to the bottom.

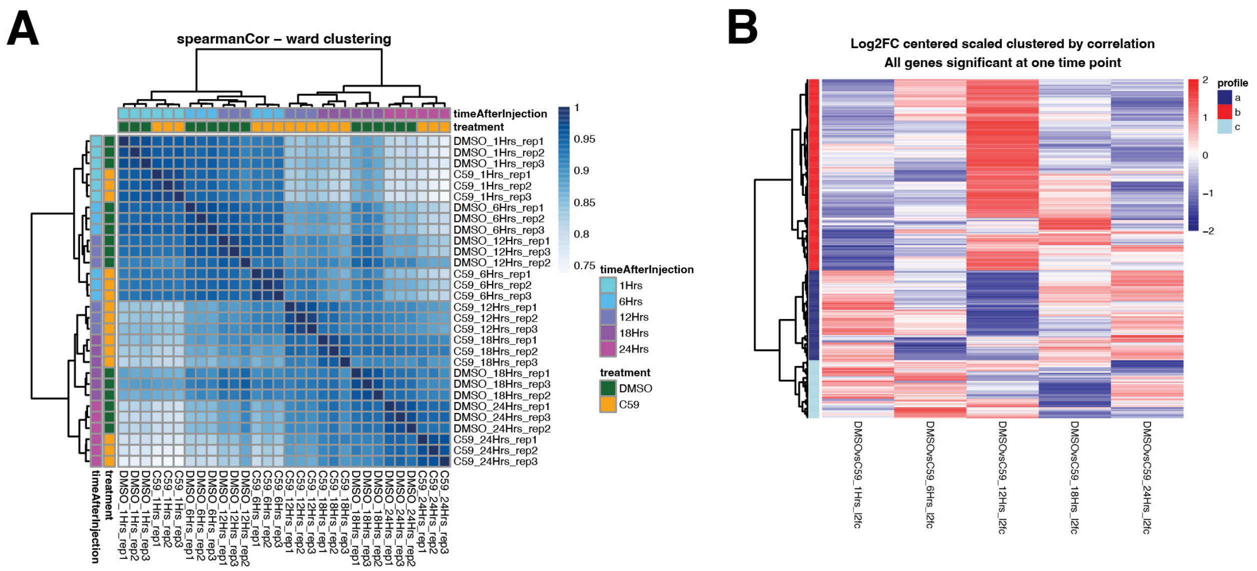


**Fig. S3. Assessment of three established small molecule inhibitors identifies C59 as the most efficacious WNT inhibitor for IP injection into pregnant mice.** (A) IP injection of a single dose of 10µg per gram body weight of either IWP2, ETC-159 and Wnt-C59 at day E10.5. The alterations of embryonic and in particular forelimb development were assessed by morphological scoring. This analysis shows that only one dose of C59 causes variable oligodactylies at high frequencies (see table below). (B) Dose-response analysis using 3 different concentrations established 10µg C59 per gram body weight as the minimal concentration that causes high frequency oligodactylies (2-4 digits, see table below). Scale bars in panels A and B are 1mm. (C) Skeletal preparations of embryos four days after DMSO or C59 IP injection at E10.5. C59-injection causes forelimb oligodactylies with three digits, while 4, 2 or only one digit are observed at much lower frequencies (see the fractions indicated). Limbs are oriented with anterior to the top and posterior to the bottom. Scale bar: 500µm.

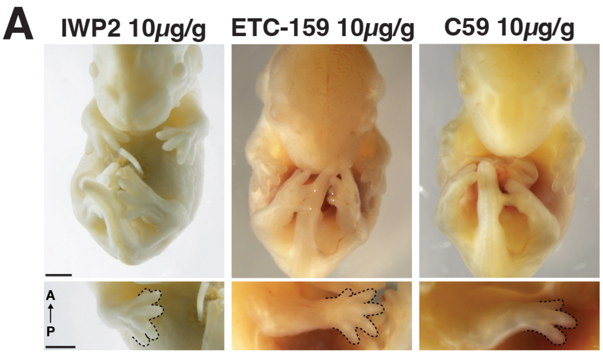


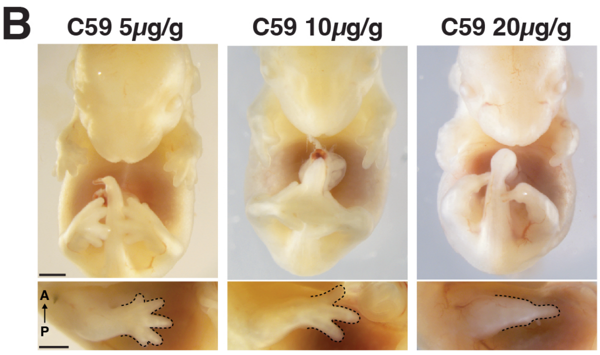


| | Right forelimb | | | |
|---|---|---|---|---|
| | control (n=54) | IWP2 10µg/g (n=35) | ETC-159 10µg/g (n=43) | C59 10µg/g (n=59) |
| 5 digits | 54 | 35 | 39 | 1 |
| 4 digits | | | 3 | 6 |
| 3 digits | | | | 43 |
| 2 digits | | | | 6 |
| 1 digits | | | 1 | 2 |
| 0 digits | | | | 1 |

| | Right forelimb | | | |
|---|---|---|---|---|
| | control (n=54) | C59 5µg/g (n=29) | C59 10µg/g (n=59) | C59 20µg/g (n=27) |
| 5 digits | 54 | 16 | 1 | 0 |
| 4 digits | | 4 | 6 | |
| 3 digits | | 6 | 43 | |
| 2 digits | | | 6 | 1 |
| 1 digits | | 3 | 2 | 25 |
| 0 digits | | | 1 | 1 |

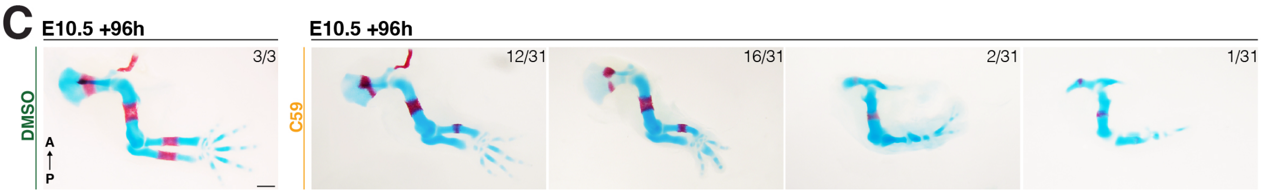


**Fig. S4. Time course RNA-seq analysis of control and C59 treated forelimb buds.** Timepoints analysed include 1, 6, 12, 18, 24 hrs after IP injection. (A) correlation plot shows the correlation between the different treatment conditions and replicates (n=3 independent biological replicates per condition ). (B) Heatmap of all DEGs clustered according to their expression profiles over time.

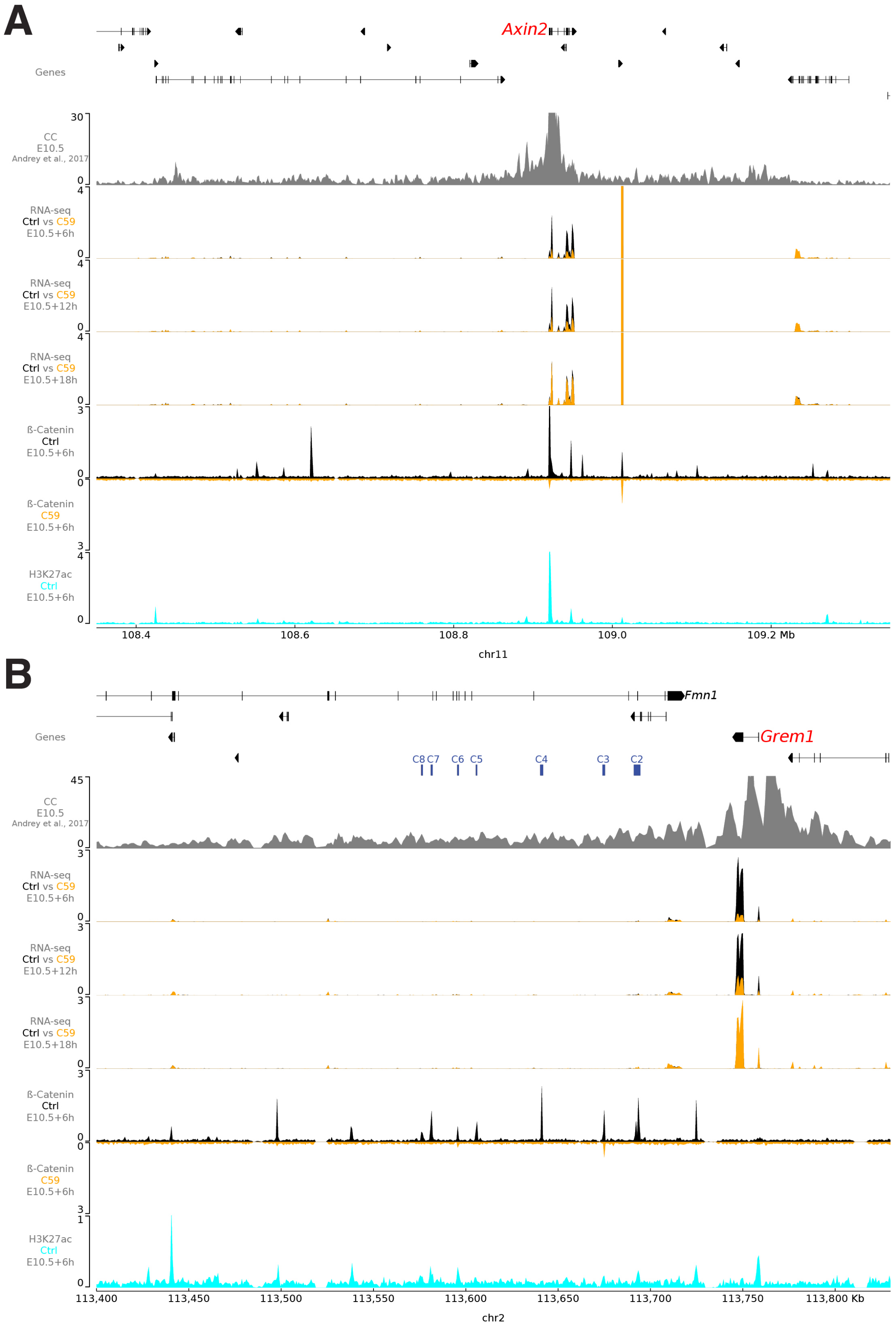

A
Axin2
Genes
CC
E10.5
Andrey et al., 2017
RNA-seq
Ctrl vs C59
E10.5+6h
RNA-seq
Ctrl vs C59
E10.5+12h
RNA-seq
Ctrl vs C59
E10.5+18h
ß-Catenin
Ctrl
E10.5+6h
ß-Catenin
C59
E10.5+6h
H3K27ac
Ctrl
E10.5+6h
108.4
108.6
108.8
109.0
109.2 Mb
chr11
B
Fmn1
Grem1
Genes
C8 C7 C6 C5 C4 C3 C2
CC
E10.5
Andrey et al., 2017
RNA-seq
Ctrl vs C59
E10.5+6h
RNA-seq
Ctrl vs C59
E10.5+12h
RNA-seq
Ctrl vs C59
E10.5+18h
ß-Catenin
Ctrl
E10.5+6h
ß-Catenin
C59
E10.5+6h
H3K27ac
Ctrl
E10.5+6h
113,400
113,450
113,500
113,550
113,600
113,650
113,700
113,750
113,800 Kb
chr2

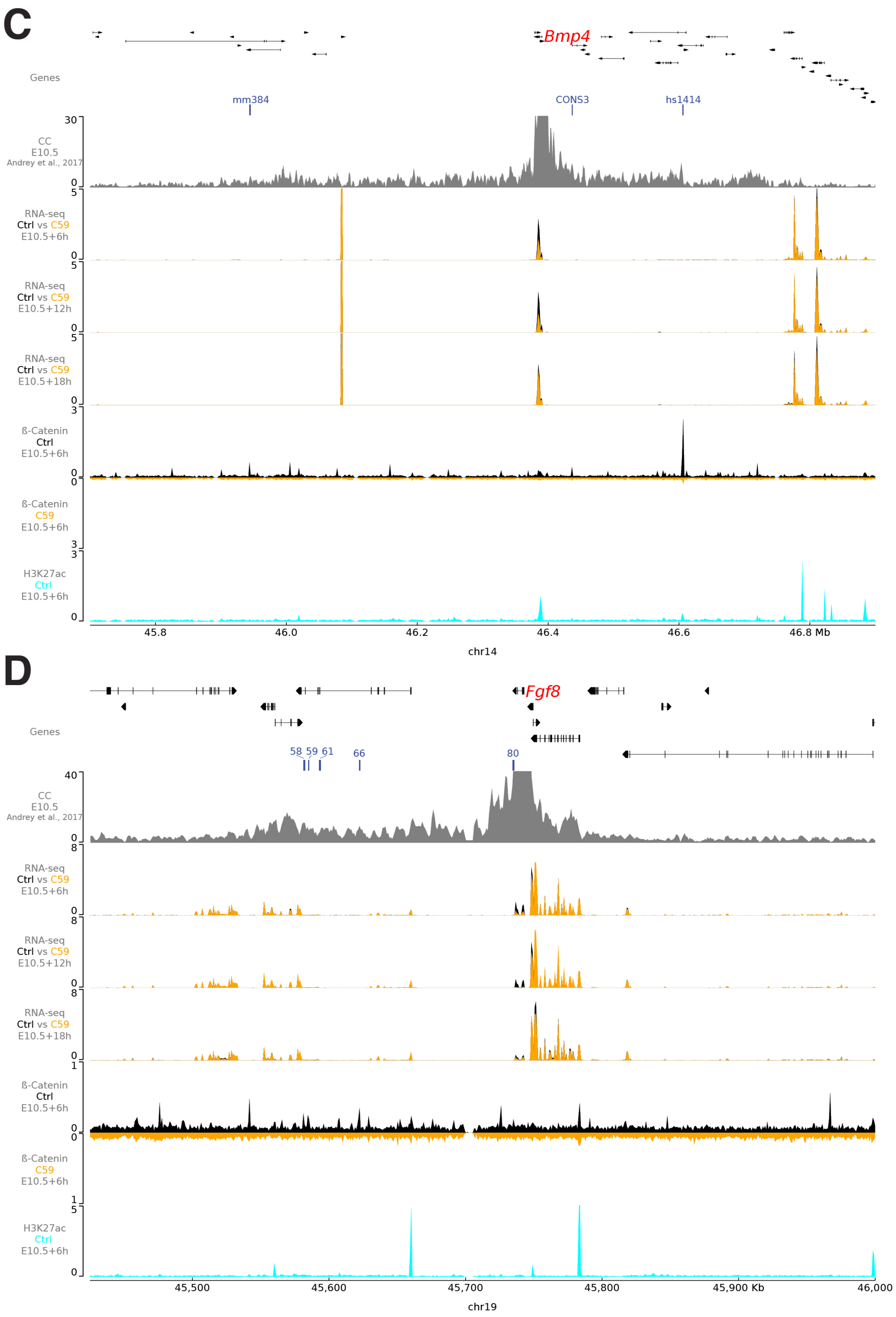
C
Bmp4
Genes
mm384
CONS3
hs1414
CC
E10.5
Andrey et al., 2017
30
0
RNA-seq
Ctrl vs C59
E10.5+6h
5
0
RNA-seq
Ctrl vs C59
E10.5+12h
5
0
RNA-seq
Ctrl vs C59
E10.5+18h
5
0
ß-Catenin
Ctrl
E10.5+6h
3
0
ß-Catenin
C59
E10.5+6h
0
3
H3K27ac
Ctrl
E10.5+6h
3
0
45.8
46.0
46.2
46.4
46.6
46.8 Mb
chr14
D
Fgf8
Genes
58 59 61
66
80
CC
E10.5
Andrey et al., 2017
40
0
RNA-seq
Ctrl vs C59
E10.5+6h
8
0
RNA-seq
Ctrl vs C59
E10.5+12h
8
0
RNA-seq
Ctrl vs C59
E10.5+18h
8
0
ß-Catenin
Ctrl
E10.5+6h
1
0
ß-Catenin
C59
E10.5+6h
0
1
H3K27ac
Ctrl
E10.5+6h
5
0
45,500
45,600
45,700
45,800
45,900 Kb
46,000
chr19

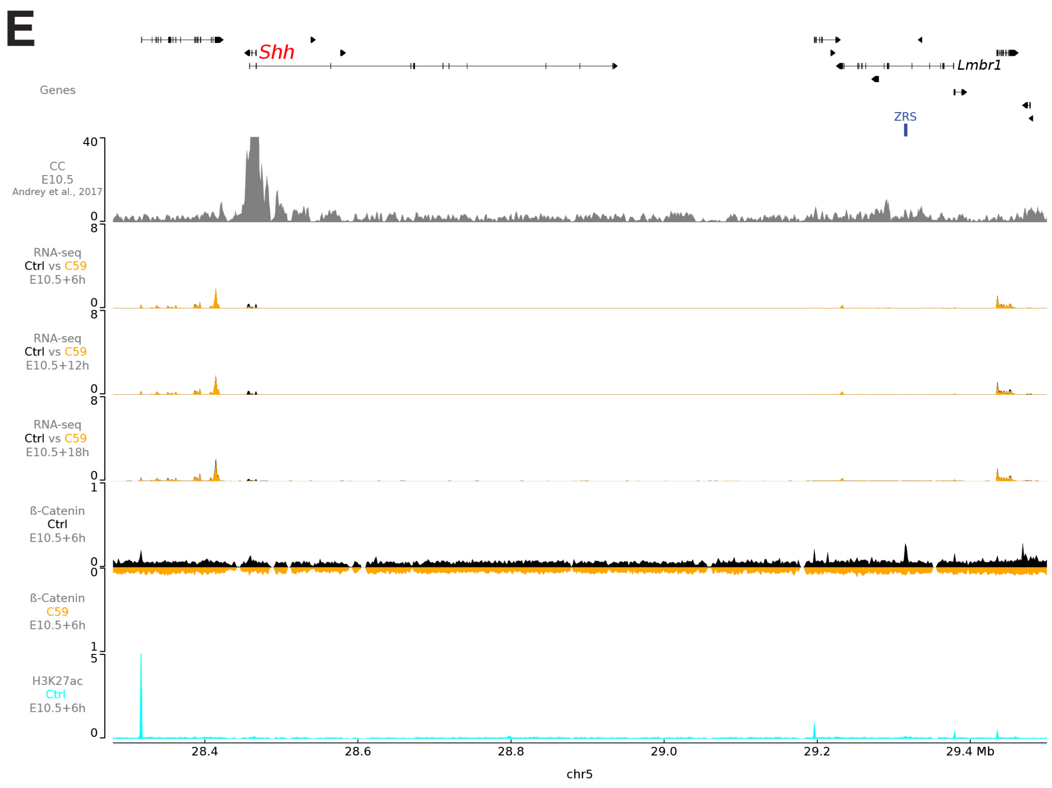


**Fig. S5. C59 disrupts β-Catenin interactions with CRM enhancers that regulate DEGs following IP injection at ~E10.5.** These graphs show additional information for the *Axin2*, *Grem1*, *Bmp4* and *Fgf8* gene regulatory landscapes (Figure 2E-H) and the *Shh* cis-regulatory landscape is included in panel E. (A-E) The regions that likely function as *cis*-regulatory landscapes for the DEGs of interest were approximated by plotting promoter-capture profiles (tracks CC, E10.5). In addition, RNAseq profiles of C59-treated (orange) forelimb buds were plotted over the control profiles (black) to reveal the alterations in expression levels at E10.5+6, E10.5+12 and E10.5+18 hours. Below the binding of β-Catenin chromatin complexes to the relevant genomic regions is shown for control (black) and C59-treated (orange, inverse) in forelimb buds at E10.5+6hrs. Together this analysis indicates that the depletion of β-Catenin chromatin complexes from known and candidate CRMs results in rapid transcriptional alteration of the associated gene (indicated in red). The histone modification H3K27Ac marks indicative for active chromatin/enhancers, shows that most β-Catenin ChIP-seq peaks are located in active chromatin regions.

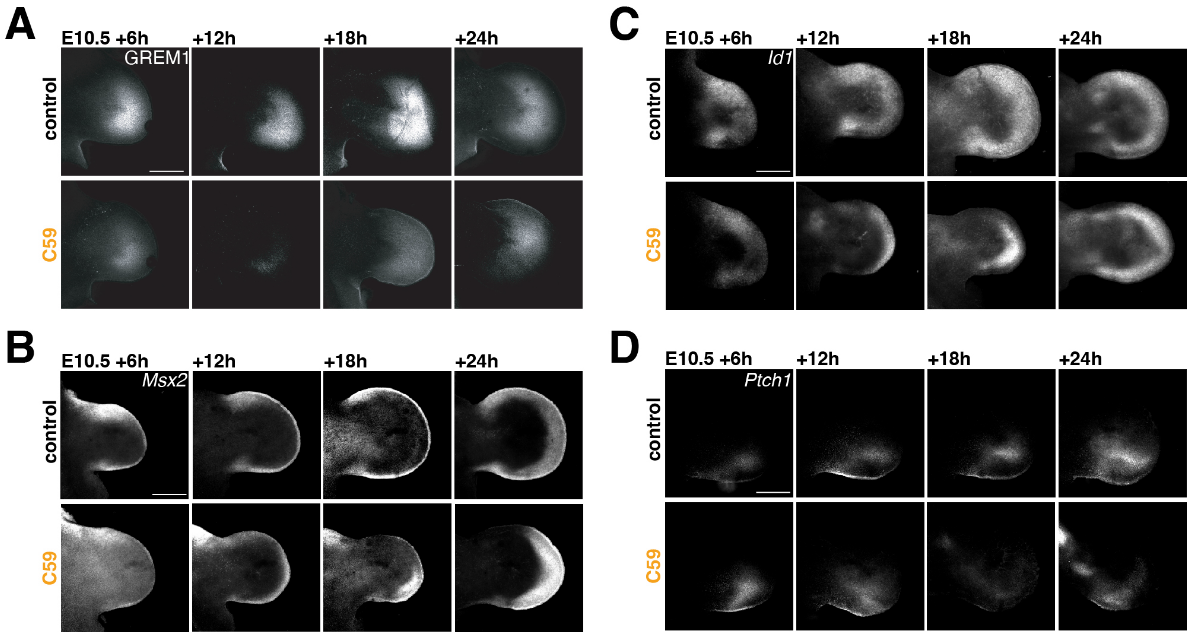


**Fig. S6. Downregulation and reestablishment of BMP and SHH target genes following C59-mediated inhibition of WNT signaling.** Spatiotemporal gene expression was assessed in control (upper panels) and C59-treated forelimb buds (E10.5, lower panels) during disruption (E10.5 +6, +12 hours) and recovery of WNT-signaling (E10.5+18, +24 hours). (A) Spatio-temporal distribution of the GREM1-protein was by whole mount immunofluorescence. This analysis reveals the rapid turnover and re-expression of GREM1 during inhibition and restoration of WNT signaling. The temporal kinetics of GREM1 loss and restoration closely follow the alterations in Grem1 transcription (Figures 3A, 4B). (B) Rapid downregulation of the BMP target gene Msx2 (E10.5+6hrs) is followed by recovery (E10.5+18-24hrs) in C59-treated forelimb buds. (C) Expression of the BMP target gene *Id1* in response to C59-mediated WNT inhibition is reduced and distalized by E10.5+12hrs and recovers in parallel to WNT signaling. (D) Expression of the SHH transcriptional target *Ptch1* is spatially reduced by E10.5+18hrs after C59-treament. n=3 independent biological samples were analysed per gene and stage. All images are shown as maximum intensity projection of the entire Z-stack range. Forelimb buds are oriented with anterior to the top and posterior to the bottom. Scale bars 200µm.

**Figure S7**

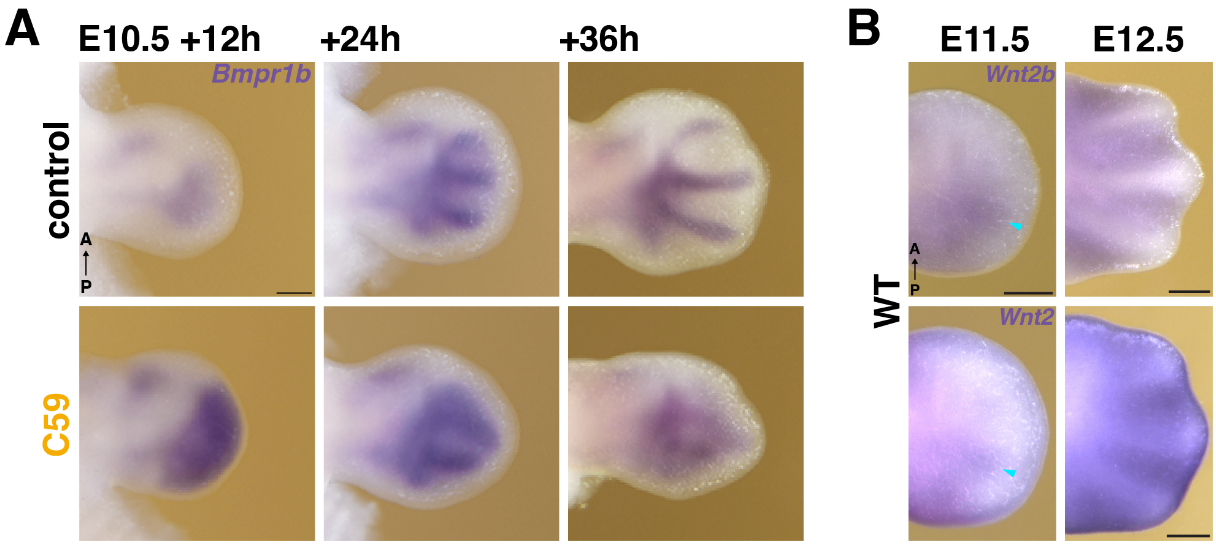


**Fig. S7. *Bmpr1b* expression dynamics during WNT signaling disruption and the spatial expression of *Wnt2* and *Wnt2b* in wildtype forelimb buds.** (A) Spatial analysis of the expression dynamics of the *Bmpr1b* receptor during disruption and recovery of WNT signaling. (B) WISH shows the *Wnt2b* (top panels) and *Wnt2* ligands (bottom) panels) expression in the presumptive interdigit mesenchyme of wildtype forelimb buds (E11.5 and E12.5). Scalebars in A and B: 250$\mu$m. A=anterior, P=posterior, D=dorsal, V=ventral. n=3 independent biological replicates were analysed for all timepoints and genes shown.

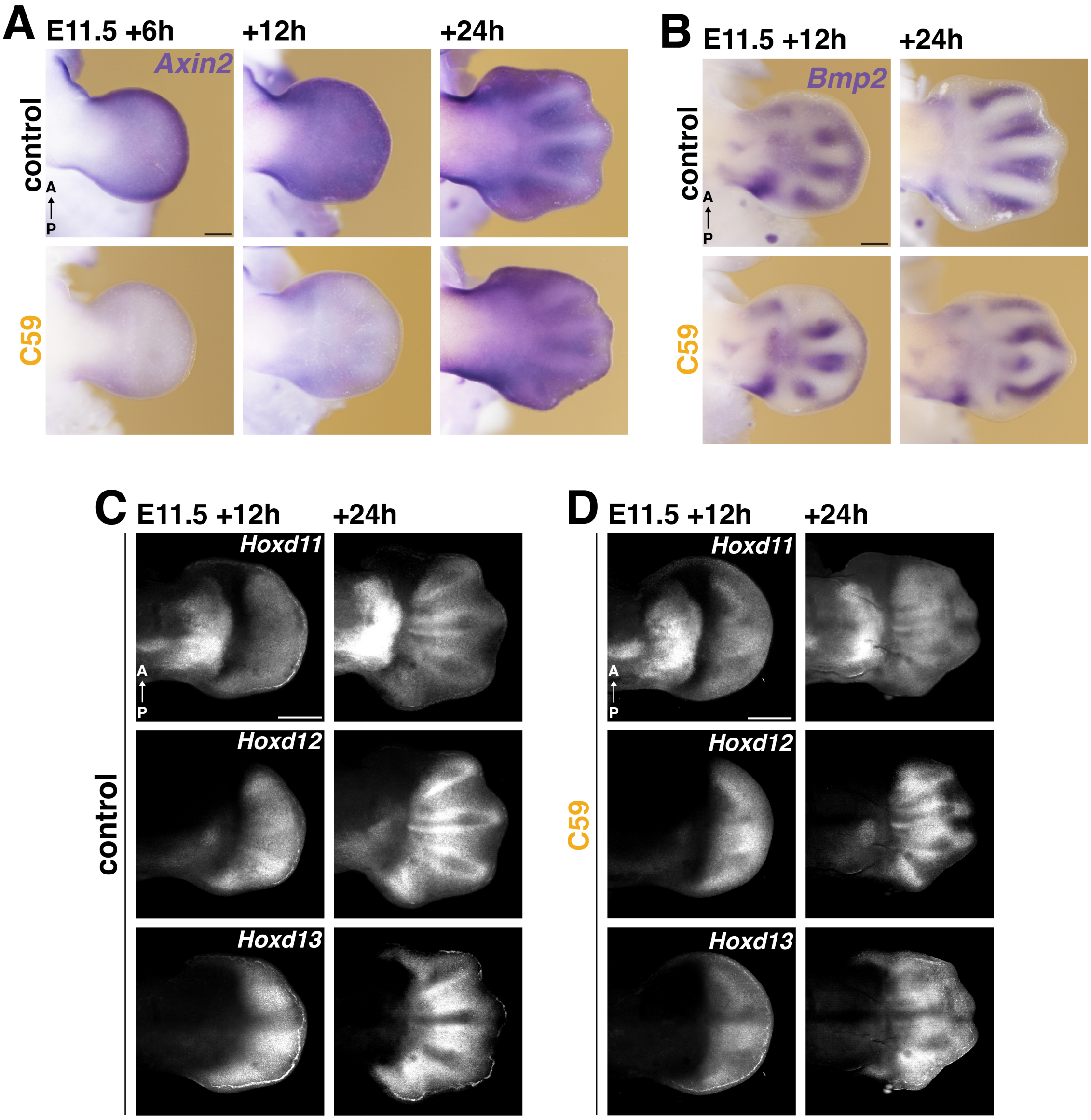


**Fig. S8. Temporal alterations and recovery of the periodic digit-interdigit patterning system after transient disruption of WNT signaling at ~E11.5** (A) *Axin2* expression in forelimb buds at E11.5+6, +12 and +24hrs following DMSO (control) and C59 treatment. This establishes that C59-mediated inhibition and recovery of WNT signaling occurs with similar temporal kinetics as at E10.5. (B) Spatio-temporal *Bmp2* expression at E11.5+12 and +24hrs after DMSO (control) and C59 treatment. (C, D) Greyscale single channels for the analysis shown in Figure 6E. *Hoxd11*, *Hoxd12* and *Hoxd13* expression in control (panel D) and C59-treated forelimb buds following C59 injection (panel E). n=3 independent biological replicates per probe and stage were analysed. All limb buds are shown with anterior to the top and posterior to the bottom. Scale bars in panel A-C: 250µm. Scale bars in panel D, E: 200µm.

**Supplementary Tables**

**Table S1. Temporal profile of differentially expressed genes.** DEGs identified at 1, 6, 12, 18 and 24hrs after C59 injection. Linked to Fig. 1, 2A.

**Table S2. DEG fold-change expression profiles over time.** DEGs were split into three clusters based on their temporal fold-change expression profiles. Linked to Fig. 1D.

**Table S3.** DEGs selected for line plots analysis of the relevant pathways. Linked to Fig 1E.

**Table S4. DEGs at the E10.5+6hrs timepoint selected for network analysis.** Linked to Fig. 2A.
